# High-Parameter Spatial Multi-Omics through Histology-Anchored Integration

**DOI:** 10.1101/2025.02.23.639721

**Authors:** Yonghao Liu, Chuyao Wang, Zhikang Wang, Liang Chen, Zhi Li, Jiangning Song, Qi Zou, Rui Gao, Bin-Zhi Qian, Xiaoyue Feng, Renchu Guan, Zhiyuan Yuan

## Abstract

Spatial omics face challenges in achieving high-parameter, multi-omics co-profiling. Serial-section profiling of complementary panels mitigates technical trade-offs but introduces the spatial diagonal integration problem. To address this, we present SpatialEx and its extension SpatialEx+, computational frameworks leveraging histology as a universal anchor to integrate spatial molecular data across tissue sections. SpatialEx combines a pre-trained H&E foundation model with hypergraph and contrastive learning to predict single-cell omics from histology, encoding multi-neighborhood spatial dependencies and global tissue context. SpatialEx+ further introduces an omics cycle module that encourages cross-omics consistency via slice-invariant mappings, enabling seamless integration without co-measured training data. Extensive validations show superior H&E-to-omics prediction, panel diagonal integration, and omics diagonal integration across various biological scenarios. The frameworks scale to datasets exceeding one million cells, maintain robustness with non-overlapping or heterogeneous sections, and support unlimited omics layers in principle. Our work makes multi-modal spatial profiling broadly accessible.

## 1. Introduction

The functionality of biological systems is intrinsically linked to the spatial organization of their cells^1^. Recent advances in spatial omics now enable high-resolution, *in situ* profiling of gene expression, epigenomic features, and proteomic distributions^2–4^. These technologies have revolutionized our capacity to map molecular architectures, offering valuable insights into tissue development^5,6^, disease progression^7,8^, and the cellular interactions^9,10^.

Although significant progress has been made, biological complexity requires a multi-layered analysis that goes beyond single-omics measurements^11^. Efforts to profile multiple omics layers from a single tissue slice— using technologies such as DBiT-seq^12^, spatial-CITE-seq^13^, spatial multimodal analysis (SMA)^14^, spatial ATAC-RNA-seq^15^, and spatial-Mux-seq^16^—face inherent trade-offs, including limited resolution, low throughput, and technical challenges due to complex processing^17,18^. An alternative approach—profiling distinct single-omics layers (e.g., transcriptomics or metabolomics) on different slices and then aligning them computationally—avoids some pitfalls, but inevitably creates mismatches at cellular level, compromising integration accuracy and downstream analysis^19–21^.

Even within a single spatial omics modality, high-parameter profiling remains challenging^3,22,23^. High-resolution spatially resolved transcriptomics (SRT) techniques, such as MERFISH^24^ and 10x Xenium^25^, achieve single-cell resolution but are limited by pre-defined gene panels, which cover only a fraction of the whole transcriptome. This limitation hinders the detection of subtle cell states and restricts the potential for hypothesis-free discovery^26–28^. Although computational imputation methods using scRNA-seq data have been proposed, they often overlook spatial context and therefore fail to capture local gene expression gradients^29^. Moreover, expanding coverage by profiling different slices with complementary gene panels brings back alignment issues similar to those seen in multi-omics workflows.

To address these limitations, we employed an experimental-computational framework that leverages serial tissue slices analyzed with distinct single-omics techniques. By pairing routine spatial single-omics assays with histology images, our goal is to reconstruct high-parameter multi-omics profiles without the need for simultaneous experimental co-profiling. However, integrating disparate omics layers across non-overlapping slices remains a significant challenge, similar to the diagonal integration issues encountered in single-cell genomics where datasets lack shared omics features^30,31^. In contrast to single-cell data, spatial omics includes additional layers of spatial context and cellular morphology that correlate with molecular profiles^32–34^. Building on these inherent correlations, we introduce the first computational framework for spatial diagonal integration, using histology as a feasible but effective anchor to align complementary omics features from adjacent slides.

We introduce SpatialEx and SpatialEx+. SpatialEx is built on a Hematoxylin and Eosin (H&E) foundation model, hypergraph learning, and a contrastive learning strategy, and it is designed to predict omics profiles directly from H&E images regardless of the spatial resolution of technologies (e.g., single-cell or spot). Building on SpatialEx, SpatialEx+ addresses the spatially resolved diagonal integration problem by connecting multiple SpatialEx models trained on adjacent slices via our newly introduced omics cycle module, which further regularizes the relationships across different omics and gene panels across slices.

Comprehensive benchmarking analyses demonstrate the superior performance of our proposed methods in a variety of tasks. These include predicting single-cell omics profiles from H&E images (i.e., H&E-to-omics prediction), using diagonal integration to expand high-parameter gene panels (i.e., panel diagonal integration), and using diagonal integration to achieve spatial multi-omics (i.e., omics diagonal integration). SpatialEx outperformed existing methods in H&E-to-omics prediction, successfully enabling the characterization of tumor microenvironments outside the sequencing area. In panel diagonal integration, SpatialEx+ successfully aligned spatial domains between two breast cancer slices that underwent SRT with non-overlapping gene panels. Furthermore, the integrated large gene panel achieved through diagonal integration revealed previously indistinguishable distinctions between immune and stromal regions. We evaluated SpatialEx+’s omics diagonal integration capabilities using two distinct cases: spatial transcriptomic-protein profiling and spatial metabolic-transcriptomic profiling. Notably, in the Parkinson’s Disease study, the SpatialEx+ generated co-profiling revealed both anatomic and fine-grained pathological domains that were overlooked by spatial single-omics alone. To further validate our approach, we tested SpatialEx+’s scalability using datasets containing over 1 million cells and challenging cases that violate the adjacent slices requirement (including slices with significant differences and no overlapping regions). In principle, SpatialEx+ enables the simultaneous generation of comprehensive high-parameter spatial multi-omics without limitation on the number of omics layers. Most significantly, SpatialEx+ achieves these capabilities while requiring only spatial single-omics assays paired with histology, thereby eliminating the need for complex paired multi-omics data collection.

## 2. Results

### The spatial diagonal integration problem

In single-cell genomics, data integration strategies are traditionally categorized into four types: vertical, horizontal, mosaic, and diagonal integration^30^. Among these, diagonal integration presents the most significant challenges due to the absence of overlapping omics features between datasets^35^. Current approaches to diagonal integration rely on weak biological links, such as correlating scATAC-seq data with gene expression through chromatin accessibility patterns around genes, or connecting genes and proteins through protein-coding gene relationships^31,36,37^. While the spatial omics field has made substantial progress in addressing vertical^38,39^, horizontal^40,41^, and mosaic integration^42^ challenges, the problem of spatial diagonal integration— specifically, how to integrate tissue slices profiled with different single-omics measurements to generate comprehensive multi-omics profiles for each cell—remains largely unresolved.

Recent work has expanded the concept of diagonal integration to include the integration of datasets with the same omics type but non-overlapping features^35^. Building on this framework, we define two scenarios in spatial diagonal integration: panel diagonal integration and omics diagonal integration. Panel diagonal integration focuses on integrating data across different slices analyzed with the same omics technology but different feature panels, while omics diagonal integration addresses the integration of different omics types across different slices. These integration approaches respectively address the challenges of achieving high-parameter (i.e., large panel) spatial profiling and spatial multi-omics profiling.

In the approach to spatial diagonal integration, one can leverage histology images as an anchoring mechanism to bridge different tissue slices. Consider a series of tissue sections, each analyzed with a distinct omics technique paired with H&E image—a much more practical approach than simultaneous spatial multi-omics co-profiling. As discussed in a recent Comment by Prof. Mingyao Li’s Lab^43^, one might initially consider developing individual H&E-to-omics prediction models for each slice, then applying these trained models across slices to predict missing omics data. This consideration led us to develop a technology-agnostic H&E-to-omics prediction model that incorporates both histology and spatial information (SpatialEx’s motivation). However, this straightforward approach proves suboptimal, as histological and spatial information alone may not provide sufficient context for accurate omics profile prediction indicated by recent studies^34,44^. This limitation motivated us to develop additional regularization modules that enable collaborative training among omics-specific H&E-to-omics models, thereby enhancing prediction accuracy using omics-omics relationships (SpatialEx+’s motivation).

### Overview of SpatialEx and SpatialEx+

Building upon our histology-anchored spatial diagonal integration framework, we first developed SpatialEx, a technology-agnostic H&E-to-omics prediction model. SpatialEx establishes a robust mapping between H&E images and omics expression profiles, enabling precise prediction of omics expression values from histological data (Fig. 1A-D). The model processes input H&E images through cell segmentation^45^ and cropping to generate single-cell morphology images, which are then transformed into high-dimensional feature representations using a pre-trained H&E foundation model^46^ (Fig. 1B).

**Fig. 1:**
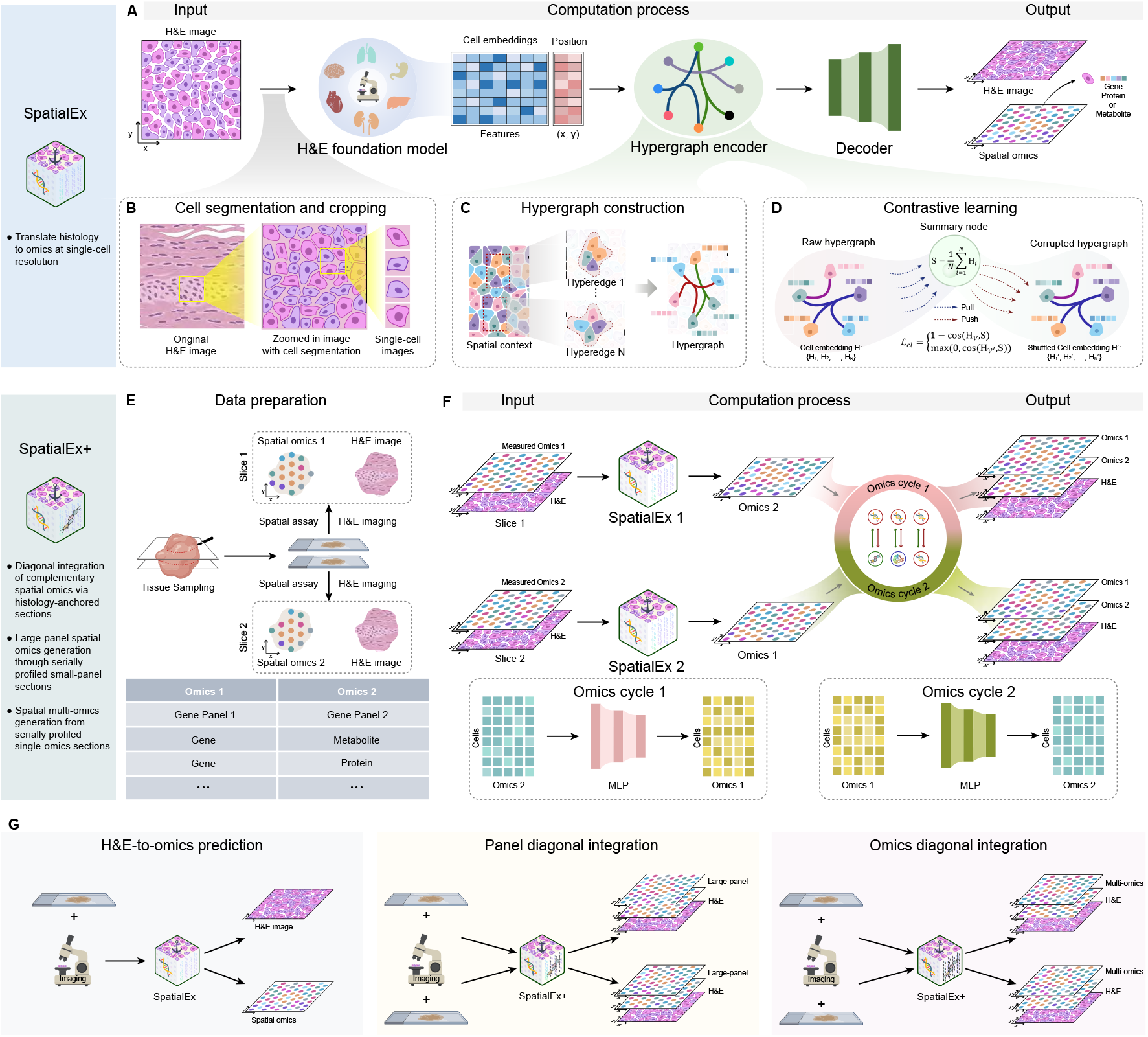
Overall architecture of SpatialEx and SpatialEx+. **A**. SpatialEx first crops image patch for each cell, which is fed into an H&E foundation model to generate low-dimensional embeddings. A cellular hypergraph is then constructed based on spatial locations, and a hypergraph encoder is applied to refines cellular features through contrastive learning. Finally, the features are decoded to predict omics profiles. **B**. The single-cell images are square patches centered on cell locations. When the cell locations are unavailable, cell segmentation is performed first. **C**. In the hypergraph, cells serve as nodes, and each cell with its spatial neighborhood forms a hyperedge. **D**. A corrupted hypergraph with feature H’ is generated by shuffling the cell embeddings H, while preserving the topology in the raw hypergraph. Contrastive learning then minimizes the discrepancy between H and the summary node S, and vice versa for the corrupted cell embeddings H’ and S. **E**. Data preparation for SpatialEx+: two slices with H&E images, each measuring different spatial omics. **F**. Building upon SpatialEx, SpatialEx+ introduces omics cycle modules to enable diagonal integration and complement the missing omics profiles in each tissue slice. **G**. Three key applications of this study include: (1) H&E-to-omics prediction, (2) Panel diagonal integration, (3) Omics diagonal integration.

While existing approaches typically employ traditional graph structures based on cell-cell adjacency to capture pairwise relationships^47–49^, a few studies suggested that such approaches represent the complexity of biological tissues in a sub-optimal manner^50,51^. To address this limitation, we implemented a hypergraph structure that models group-level correlations through hyperedges (Fig. 1C), motivated by established principles in spatial biology, i.e., cellular microenvironments require collective interactions among cell groups to regulate omics states^52^.

To further strengthen the model’s capabilities, we incorporated contrastive learning to integrate globally relevant data across the entire tissue (Fig. 1D). This approach involves generating a corrupted hypergraph H’ by shuffling cell embeddings while maintaining the topological structure. The contrastive learning mechanism ensures that cells in the original hypergraph are drawn closer to the summary node S while pushing cells in the corrupted hypergraph farther away, thereby effectively capturing both cell expression patterns and their surrounding microenvironments. During testing, while the contrastive learning component is omitted, all other procedures remain operational.

Building on SpatialEx, we developed SpatialEx+ to enable diagonal integration of spatial omics data from adjacent tissue slices (Fig. 1E, F). The framework begins with data preparation (Fig. 1E), where spatial assay and H&E staining are performed on adjacent slices of a tissue sample, generating both spatial omics data and H&E images for various molecular modalities including genes, metabolites, or proteins. SpatialEx+ implements a collaborative training framework that enables mutual guidance between predictive branches of different omics types. When processing H&E images from adjacent tissue slices, the framework simultaneously generates distinct omics data for both slices. Central to SpatialEx+ is the *omics cycle module*, which employs a non-linear transformation to establish explicit omics-omics associations. This module encourages conservation of cross-omics mapping functions between adjacent tissue sections: the relationship *f* (e.g., gene expression → protein) remains identical between slices, even when omics 1 is measured/predicted in Slice 1 and omics 2 is predicted/measured in Slice 2. By constraining *f* to align across neighboring sections—where spatial continuity ensures biological similarity—the model avoids conflicting slice-specific predictions, harmonizing multi-omics inferences under unified tissue-wide principles.

We validated our framework across three experimental scenarios (Fig. 1G): (1) H&E-to-omics prediction, where SpatialEx translates histology to omics at single-cell resolution; (2) panel diagonal integration, where SpatialEx+ generates large-panel SRT data containing both gene panels from adjacent slices; and (3) omics diagonal integration, where SpatialEx+ generates spatial multi-omics data containing both omics from adjacent slices. Detailed procedures can be found in Extended Data Fig. 1, Supplementary Text 1 and Supplementary Fig. 1.

### SpatialEx translates histology to omics at single-cell resolution

Current H&E-to-omics prediction methods are mostly limited to spot-level resolution^32,53–55^, leaving a critical gap in single-cell resolution prediction. While DeepPT^55^ represents the most recent advance and state-of-the-art in this field based on a recent benchmarking study^34^, it was originally designed for spot-resolution prediction. We modified DeepPT’s implementation to enable single-cell predictions, creating the first baseline for comparison at this resolution. In addition, we also introduced a CNN-based regression model (CNN_Reg) for comparison. CNN_Reg uses ImageNet-pretrained ResNet50 to extract image representations, followed by a multi-layer perceptron to learn the mapping from H&E features to gene expression. Then, we adapted Hist2ST^53^ and THItogene^56^ for single-cell data by employing a mini-batch training strategy. In contrast, SpatialEx was designed as a resolution-agnostic framework, capable of accurate predictions at both spot and single-cell levels without modification. Using formalin-fixed paraffin-embedded (FFPE) sections from a 10x Xenium Human Breast Cancer dataset^25^, along with one normal tissue (human colon), one diseased tissue (human skin), and one mouse tissue (mouse colon), we evaluated SpatialEx’s single-cell prediction capability through a train-test framework (Fig. 2A, B). Both training (Slice 1) and test (Slice 2) sections underwent Xenium sequencing for spatially resolved gene expression profiling and H&E staining for histology.

**Fig. 2:**
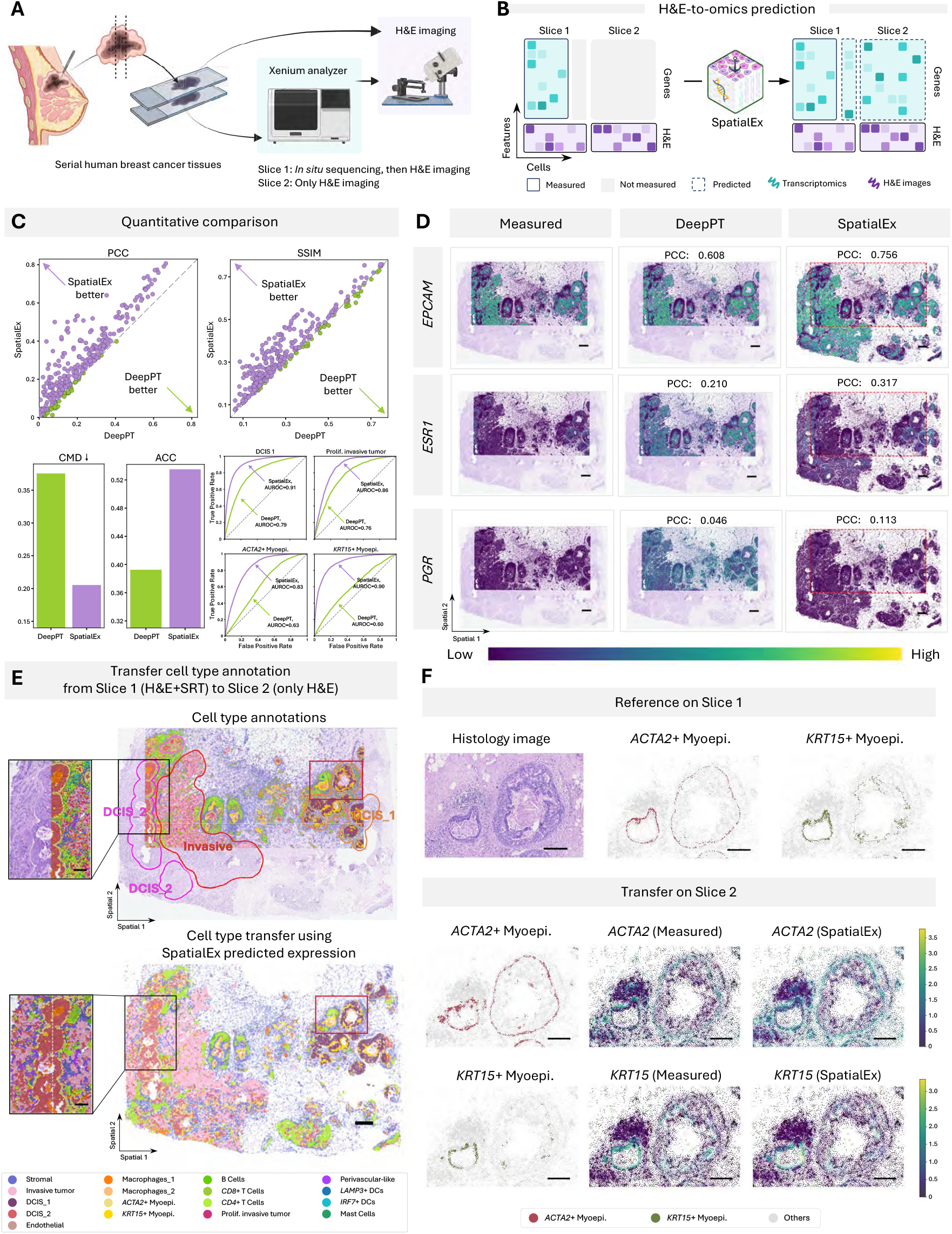
SpatialEx translates histology to omics at single-cell resolution. **A**. Experimental process: serial human breast cancer tissues were imaged, with *in situ* sequencing only performed on Slice 1. **B**. Computational schematic: SpatialEx links H&E features with single-cell omics within the sequencing area of Slice 1, then infers single-cell omics profiles across the whole slide images of both slices. **C**. Top, from left to right: scatter plots of PCC, SSIM. Each dot represents one of the 313 genes. Bottom, from left to right: bar plot of CMD; bar plot of the cell type transfer ACC using predicted gene expression; ROC curves of four cell types, with AUROC annotated. Dashed lines indicate random guess. **D**. Gene expression of *EPCAM, ESR1*, and *PGR*. The red box indicates the sequencing area. **E**. Visualization of cell type transfer results. Top: annotations on Slice 2. Bottom: transferred results from Slice 1 to Slice 2 using SpatialEx predicted gene expressions. The zoomed-in ROI (black box) illustrates the transferred cell types outside the sequencing area. **F**. The zoomed-in ROI (red box in Fig. 2E). Top: H&E image and spatial distribution of *ACTA2+* and *KRT15+* Myoepi. on Slice 1. Bottom, from left to right: transferred cell type using SpatialEx predicted gene expression; measured expression of corresponding marker genes; SpatialEx predicted markers expression on Slice 2. (Scale bars in D, E: 500 um, in F: 250 um)

Two metrics, including Pearson Correlation Coefficient (PCC) and Structural Similarity Index Measure (SSIM), are utilized for quantitative performance evaluation, with higher PCC and SSIM values indicating better performance. Fig. 2C illustrates the quantitative comparison between SpatialEx and DeepPT in terms of prediction evaluation and downstream cell type classification, while Extended Data Fig. 2 shows the comparison of SpatialEx against CNN_Reg and DeepPT in terms of prediction evaluation across multiple tissue datasets. In the scatter plots for PCC and SSIM (Fig. 2C), each dot represents an individual gene. Notably, compared to the two baseline models, the proposed SpatialEx demonstrates significantly improved performance in predicting the gene expression. A more comprehensive quantitative and qualitative comparison across all baseline methods is presented in Extended Data Fig. 3.

In Fig. 2D, we visualized the gene expressions of *EPCAM, ESR1*, and *PGR* predicted by SpatialEx and measured by spatial assays (More visualizations of genes can be found in Extended Data Fig. 4). Notably, *ESR1* and *PGR* are genes encoding biomarkers for breast cancer prognosis^25,33^. Based on the comparison, SpatialEx demonstrates a significantly better prediction capability. The results from DeepPT’s exhibit either under-expressions or over-expressions, deviating significantly from the real patterns. Moreover, SpatialEx not only effectively predicts the sequenced regions within the red box, with its predicted gene expression closely matching the measured gene expression, but also extends predictions to areas with only H&E available (outside the red box).

To evaluate whether the predicted gene expressions retain biological relevance, we first computed the correlation matrix distance (CMD) metric (Fig. 2C and Extended Data Fig. 2A), where SpatialEx achieved a score of 0.206, substantially lower than that of DeepPT (0.302) and CNN_Reg (0.271), indicating better agreement with the measured data (lower is better). We further evaluated the correlation between the spatial autocorrelation (Moran’s *I*) of the measured and predicted genes, demonstrating that SpatialEx more effectively preserves spatial autocorrelation patterns (Supplementary Fig. 2). Meanwhile, spatial domain identification using CellCharter^40^ on the hColon_Non_diseased, mouse_Colon, and hSkin_Melanoma datasets revealed that domains derived from SpatialEx-predicted gene expression closely resembled those obtained using the measured gene expression across various spatial resolutions (Extended Data Fig. 5).

To further assess biological relevance at single-cell resolution, we evaluated cell type annotation accuracy (ACC) on the Human breast cancer dataset using predicted gene expression profiles. SpatialEx demonstrated superior performance across four major cell types in Receiver Operating Characteristic (ROC) (Fig. 2C), with comprehensive evaluation of additional cell types provided in Extended Data Fig. 6A. The spatial distribution of cell types annotated using SpatialEx’s predicted gene expression (Fig. 2E bottom) showed strong concordance with annotations based on measured gene expression in Slice 2 (Fig. 2E top). In contrast, the cell type annotations derived from DeepPT’s predicted expression appeared noticeably more disorganized (Extended Data Fig. 6B). Importantly, SpatialEx enabled cell type annotation in regions where only H&E images were available (beyond the sequenced area), which would be impossible without the SRT predicted by SpatialEx on Slice 2 (Fig. 2E). This predicted result indicates that SpatialEx can easily extend to SRT-unmeasured regions, revealing more comprehensive tissue architecture across larger tissue.

Then, we focused on two biologically distinct subtypes of myoepithelial cells characterized by marker genes: *KRT15* and *ACTA2*, within the region of interest (ROI) highlighted by the red box in Fig. 2E. These subtypes are identifiable in Slice 1 by annotation (Fig. 2F top) and, due to tissue continuity, are expected to have a similar distribution on Slice 2. As shown in Fig. 2F bottom, SpatialEx successfully predicted the expression of *KRT15* and *ACTA2* on Slice 2, enabling accurate subtype annotations. In contrast, the prediction from DeepPT is far from stratifying, failing to transfer the cell-type annotations successfully (Extended Data Fig. 6C). Looking across the entire tissue section (Fig. 2E top), in the DCIS_1 region, far from the invasive tumor areas, both *ACTA2+* and *KRT15+* myoepithelial cells are prominent. However, near the more invasive DCIS_2 areas, only *ACTA2*+ myoepithelial cells are observed. Then, in invasive tumor regions, both myoepithelial cell subtypes are absent. This finding confirms that the diminishment of myoepithelial cells is a critical hallmark of the transition of cancer from *in situ* to invasive stages^25^.

### SpatialEx+ enables larger panel spatial analysis through panel diagonal integration

High-resolution spatial omics technologies typically face inherent limitations in feature panel size, e.g., spatial transcriptomic or protein profiling^4^. Using spatial transcriptomics as an example, even advanced platforms like 10x Xenium can only profile a restricted number of genes simultaneously. We developed SpatialEx+ to overcome this fundamental constraint through panel diagonal integration (Fig. 3A). The approach combines experimental profiling of different gene panels across tissue sections with computational integration using H&E images as anchors. While we modified DeepPT to enable independent model building for each panel, SpatialEx+ broke this isolation through its novel omics cycle module, which uniquely regularizes each omics-specific prediction.

**Fig. 3:**
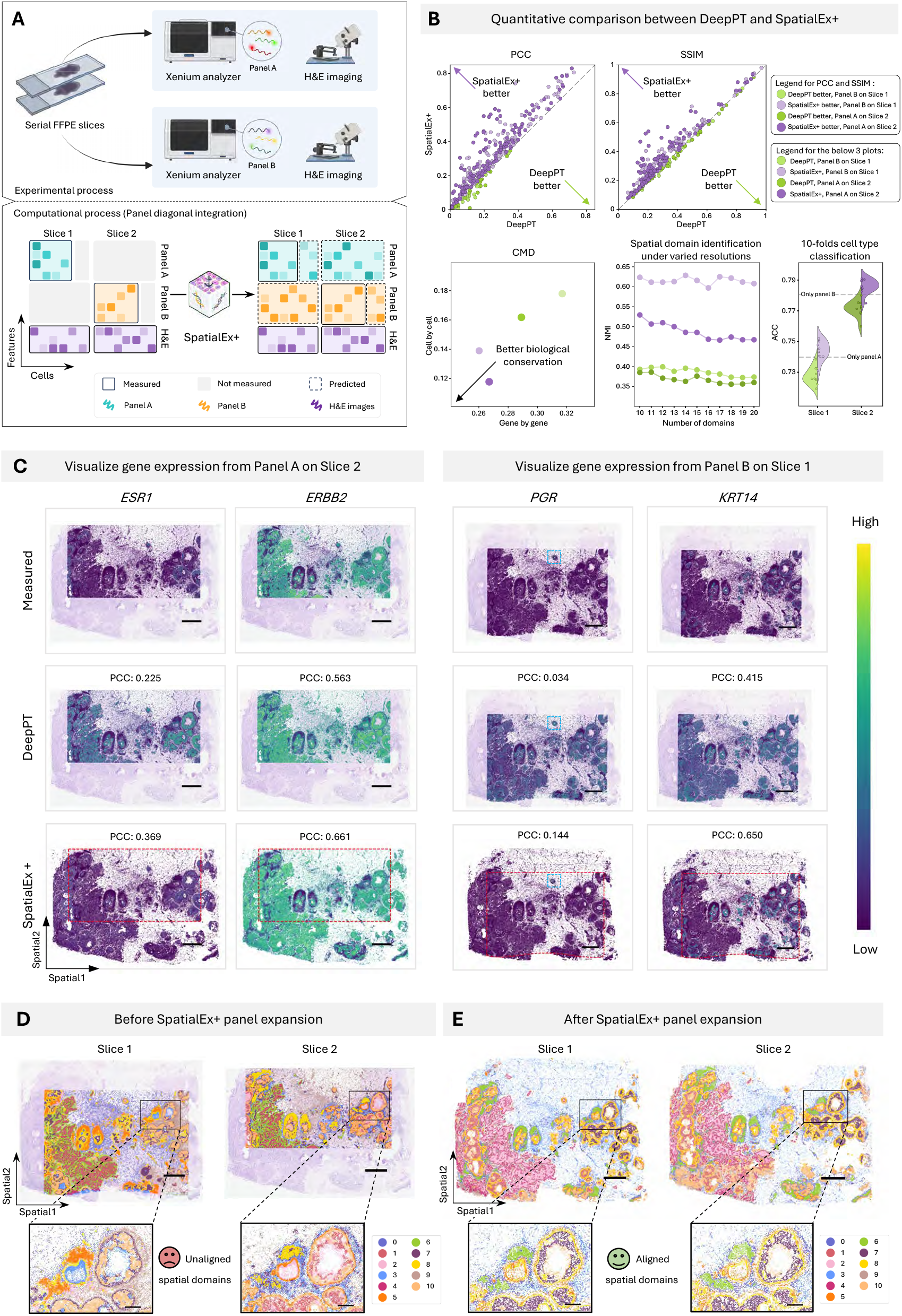
SpatialEx+ enables larger panel spatial analysis through panel diagonal integration. **A**. Schematic diagram for SpatialEx+ panel diagonal integration. For experimental process, two serial tissue sections were sequenced using Xenium, measuring Panel A on Slice 1 and Panel B on Slice 2. For computational process, SpatialEx+ bridges the diagonal gene expression matrix by H&E images, enabling more comprehensive gene expression across both slices. **B**. Quantitative comparison between SpatialEx+ and DeepPT. Top: scatter plots of PCC and SSIM. Each dot represents one of the 313 genes (150 for Panel A and 163 for Panel B). Bottom, from left to right: scatter plot of the CMD on the two slices, with the x- and y-axis representing the gene and cell dimension, respectively; line plot of NMI of spatial domain identification using predicted panels across varying domain numbers; the violin plot of ACC from 10-fold cross-validation for cell type transfer task using predicted full-panel gene expression, with the dash line indicating the ACC using the measured small-panel. **C**. Visualization of gene expressions: *ESR1* and *ERBB2* on Slice 2, and *PGR* and *KRT14* on Slice 1. The red box indicates the sequencing area. **D**. Left: spatial domain identified on Slice 1 using measured Panel A. Right: spatial domain identified on Slice 2 using measured Panel B. **E**. Spatial domain identified on the two slices using SpatialEx+ predicted full-panel (Panel A + B). (Scale bars in C, D, E: 1 mm)

To evaluate SpatialEx+’s panel expansion capabilities, we applied a complementary panel design using the 10x Xenium Human Breast Cancer dataset^25^. We divided its 313-gene panel into two non-overlapping sets (Panel A: 150 genes, Panel B: 163 genes) and assigned Panel A to Slice 1 and Panel B to Slice 2 for measurement, creating a scenario where adjacent sections have complementary gene coverage. The measured expression of the held-out panel (Panel B in Slice 1 and Panel A in Slice 2) served as ground truth for quantitative evaluation. Quantitative assessment demonstrated SpatialEx+’s superior performance across multiple metrics (Fig. 3B). Gene-level comparisons showed consistently higher PCC and SSIM values compared to DeepPT. In this experimental setting, we also validated the effectiveness of each component of SpatialEx+, as shown in Supplementary Fig. 3.

To validate biological relevance, CMD metrics confirmed SpatialEx+’s enhanced ability to preserve both cell-to-cell and gene-to-gene relationships present in the original data. In addition, SpatialEx+ more effectively maintained the spatial autocorrelation of gene expression (Supplementary Fig. 4). We further evaluated spatial domain identification using CellCharter (see Methods). SpatialEx+ achieved (28%-66%) improvement in normalized mutual information (NMI) (computed against CellCharter labels using measured full panel for each clustering resolution) across various resolution scales compared to DeepPT, demonstrating robust performance (Fig. 3B and Supplementary Fig. 5). Quantitative results of spatial domain identification across different tissue types (such as skin melanoma and healthy colon) are shown in Supplementary Fig. 6, demonstrating the generalizability of SpatialEx+ across diverse tissue types. Cell type annotation accuracy, assessed through 10-fold cross-validation, showed significant improvement when using SpatialEx+’s expanded gene panels compared to single-panel baseline (indicated by gray horizontal line in Fig. 3B). This improvement highlights both the importance of comprehensive gene panels and SpatialEx+’s effectiveness in panel expansion.

In Fig. 3C, we presented the measured expressions of the genes *ESR1* and *ERBB2* in Slice 1, as well as *PGR* and *KRT14* in Slice 2, alongside the predicted expression distributions obtained using DeepPT and SpatialEx+. Among these genes, the estrogen receptor (ER/*ESR1*), progesterone receptor (PR/*PGR*), and human epidermal growth factor receptor 2 (HER2/*ERBB2*) are three crucial ones for breast cancer subtyping^25^. Upon the results in Fig. 3C, it is evident that the gene expressions from DeepPT significantly deviate from the ground truth and are limited to the sequencing-covered regions. The DeepPT predicted *ESR1+* covers the majority of the slice, conflicting the measured results significantly. Additionally, the *PGR* gene is expressed in only a small subset of cells within the blue box, which can be successfully captured only by SpatialEx+ (Fig. 3C). This finding underscores SpatialEx+’s ability to accurately predict the expression of disease-related genes.

The impact of panel expansion became evident in spatial domain analysis. Initial domain identification using separate panels showed marked inconsistencies between slices (Fig. 3D). After SpatialEx+ integration, spatial domains showed strong alignment between sections (Fig. 3E), enabling unified tissue architecture analysis impossible with single-panel approaches (comparative analysis with DeepPT shown in Supplementary Fig. 7). This demonstrates SpatialEx+’s ability to overcome coverage limitations while maintaining biological fidelity.

### Scalability on million-cell tissue sections

To validate scalability, we applied SpatialEx+ to consecutive large-scale tissue sections from the 10x Xenium website, each containing approximately 900,000 cells (Fig. 4A). To simulate complementary panel analysis, we divided the 280-gene Xenium panel equally between the two adjacent sections and applied SpatialEx+ for diagonal integration using H&E images as anchors (Fig. 4B).

**Fig. 4:**
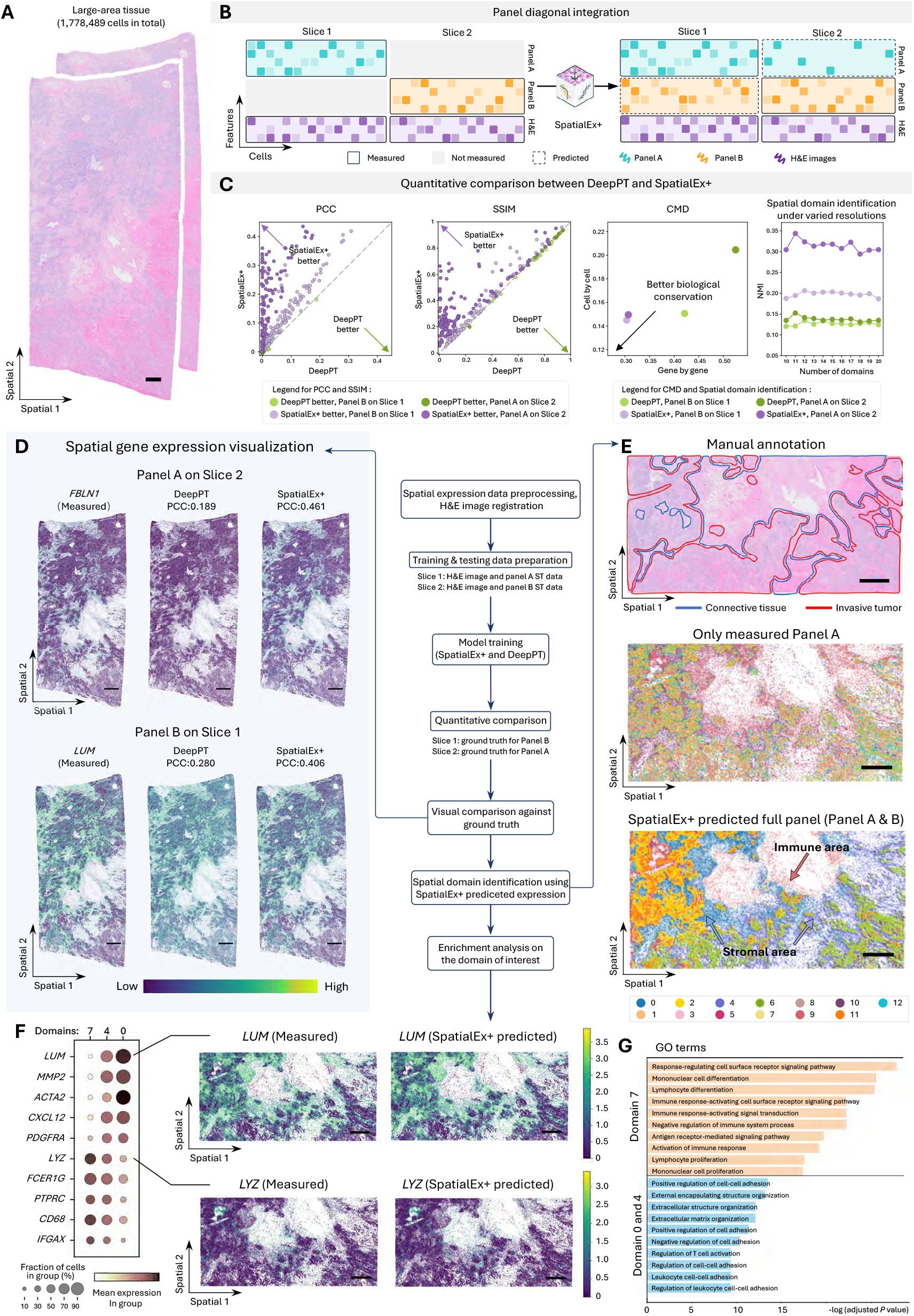
Scalability on million-cell tissue sections. **A**. The H&E image of large-area tissues from the Xenium Human Breast IDC Big dataset. **B**. The computational process for analyzing two large tissue slices. **C**. Quantitative comparison of DeepPT and SpatialEx+ performance. From left to right: scatter plots of PCC and SSIM, with each dot represents one of the 280 genes (140 for Panel A and 140 for Panel B); scatter plot of the CMD between SpatialEx+ and DeepPT on the two slices, with the x- and y-axis representing the gene and cell dimension, respectively; line plot of NMI of spatial domain identification using predicted panels across varying domain numbers. **D**. Visualization of gene expressions: *FBLN1* on Slice 1 and *LUM* on Slice 2. **E**. Top: pathologist annotated H&E images of Slice 1. Center: spatial domains identified using measured Panel A. Bottom: spatial domains identified using SpatialEx+ predicted full-panel (Panel A + B). **F**. Left: dot plot of differentially expressed genes in domains 0, 4, and 7 from Fig. 4E bottom. Right: spatial expressions of *LYZ* and *LUM* in the ROI. **G**. GO terms enriched for the markers of domain 7, and domain 0&4. The length of the bar represents the −log10 (adjusted *P* value) from one-tailed Fisher’s test. *P* values were corrected for multiple-hypothesis testing using the Benjamini–Hochberg method. (Scale bars in A, E, F: 1 mm, in D: 1.5 mm)

Quantitative evaluation demonstrated SpatialEx+’s superior performance at scale (Fig. 4C). Gene-level comparisons showed consistently higher PCC and SSIM values compared to DeepPT, while CMD measurements confirmed better preservation of both gene-gene and cell-cell relationships (Fig. 4C). Following the same spatial domain identification setup as in the previous section, SpatialEx+ again significantly outperformed DeepPT across varying numbers of domains (Fig. 4C).

We validated biological relevance through analysis of cancer-associated genes (Fig. 4D). In Fig. 4D, we visualized the gene expressions of *FBLN1* in Slice 2 and *LUM* in Slice 1, predicted by SpatialEx+ and DeepPT. *FBLN1* is associated with cell transformation and tumor invasion^57,58^, promoting breast cancer cell survival during doxorubicin treatment^59^. And previous study has shown that lumican^60^, encoded by *LUM*, is overexpressed in the breast cancer stroma and may influence tumor invasion and metastasis by modulating matrix structure and collagen arrangement. Based on the visualizations, DeepPT fails to effectively capture the spatial expression patterns of these two genes, as evidenced by numerous instances of abnormal over-expressions or under-expressions (Fig. 4D). In contrast, the gene expression predicted by SpatialEx+ closely resembles the ground truth patterns for both genes. We visualized spatial expression patterns of more genes in Extended Data Fig. 7.

The impact of panel expansion became evident in fine-grained spatial analysis (Fig. 4E). Fig. 4E illustrates the zoomed in tissue region in terms of identified spatial domains in Slice 1 across different analysis settings, including measured Panel A only (Fig. 4E middle) and measured Panel A and Panel B (Fig. 4E bottom). When only measured Panel A is used for analysis, the detected domains lead to crucial fine-grained domains being completely overlooked (Fig. 4E middle). By combining the measured genes with the predicted genes, we successfully distinguished the immune area and the stromal area (validated in the following functional analysis), as indicated by the arrows at the bottom of Fig. 4E. The domain distribution closely resembles the results obtained using the two measured full panel in Supplementary Fig. 8. Notably, we consulted senior pathologists to annotate this tissue region (Fig. 4E top), but they were unable to distinguish the immune and stromal regions based on histological features, indicating the additional insights brought by the gene expression information. We next examined these distinctions through functional analysis. Given the domains 0, 4, and 7 are particularly identified after integrating the gene panels, we performed differential gene expression analysis on these domains, and found that the critical markers for these regions, such as *LUM* and *LYZ*, are primarily located on Panel B (Fig. 4F and Supplementary Fig. 9). Gene ontology (GO) analysis of these domains (Fig. 4G) provided functional characterization of the tissue architecture: Genes highly expressed in Domain 7 were enriched for immune response pathways (GO:0002253, adjusted *P* value=4.69e-9), while Domains 0 and 4 showed enrichment for extracellular matrix and cell adhesion processes (GO:0022407, adjusted *P* value=5.61e-6). These molecular signatures from the expanded panel were essential for comprehensive characterization of the tissue’s functional organization, highlighting the necessity of panel expansion for detailed spatial analysis.

### SpatialEx+ enables spatial multi-omics through omics diagonal integration

Given the technique challenges associated with spatial multi-omics sequencing from assay, existing studies generate the spatial multi-omics data by aligning serially profiled single-omics sections to construct a multi-omics dataset. Nevertheless, variations in cellular distribution between sections render this alignment strategy suboptimal. In this section, we employed SpatialEx+ to reconstruct the spatial multi-omics data in an end-to-end function with H&E images as the linkage anchor. The omics diagonal integration analysis was performed on two datasets to demonstrate its potential and applicability.

The first transcriptomic-protein integration dataset is from Human Breast Cancer dataset^25^, which consists of two slices, with the first one sequenced by Xenium and Immunofluorescence, and the second one only sequenced by Xenium. We utilized the protein data of Slice 1 and gene expression data of Slice 2 to simulate the serially profiled sections with different omics (Fig. 5A). We presented the spatial expression of the protein HER2 and CD20 in Fig. 5B, including the measured expression on Slice 1, the SpatialEx+ expanded expression to the whole slide on Slice 1, and the whole slide on Slice 2. Additionally, we illustrated the spatial expression of their corresponding marker genes, with *ERBB2* encoding HER2 and *MS4A1* encoding CD20. The visualization demonstrates that SpatialEx+ not only accurately predicted their expression patterns but also expanded the sequencing region, ensuring a comprehensive panorama of the whole breast cancer slice. At the same time, we specifically zoomed in the spatial distribution of CD20 and its corresponding gene *MS4A1*. The remarkable consistency of expressions across different slices validates that SpatialEx+ can precisely capture fine-grained spatial omics patterns. These results demonstrate that our SpatialEx+ effectively performs omics diagonal integration while generating both omics layers for a single tissue slice. Quantitative performance metrics are detailed in Extended Data Fig. 8A.

**Fig. 5:**
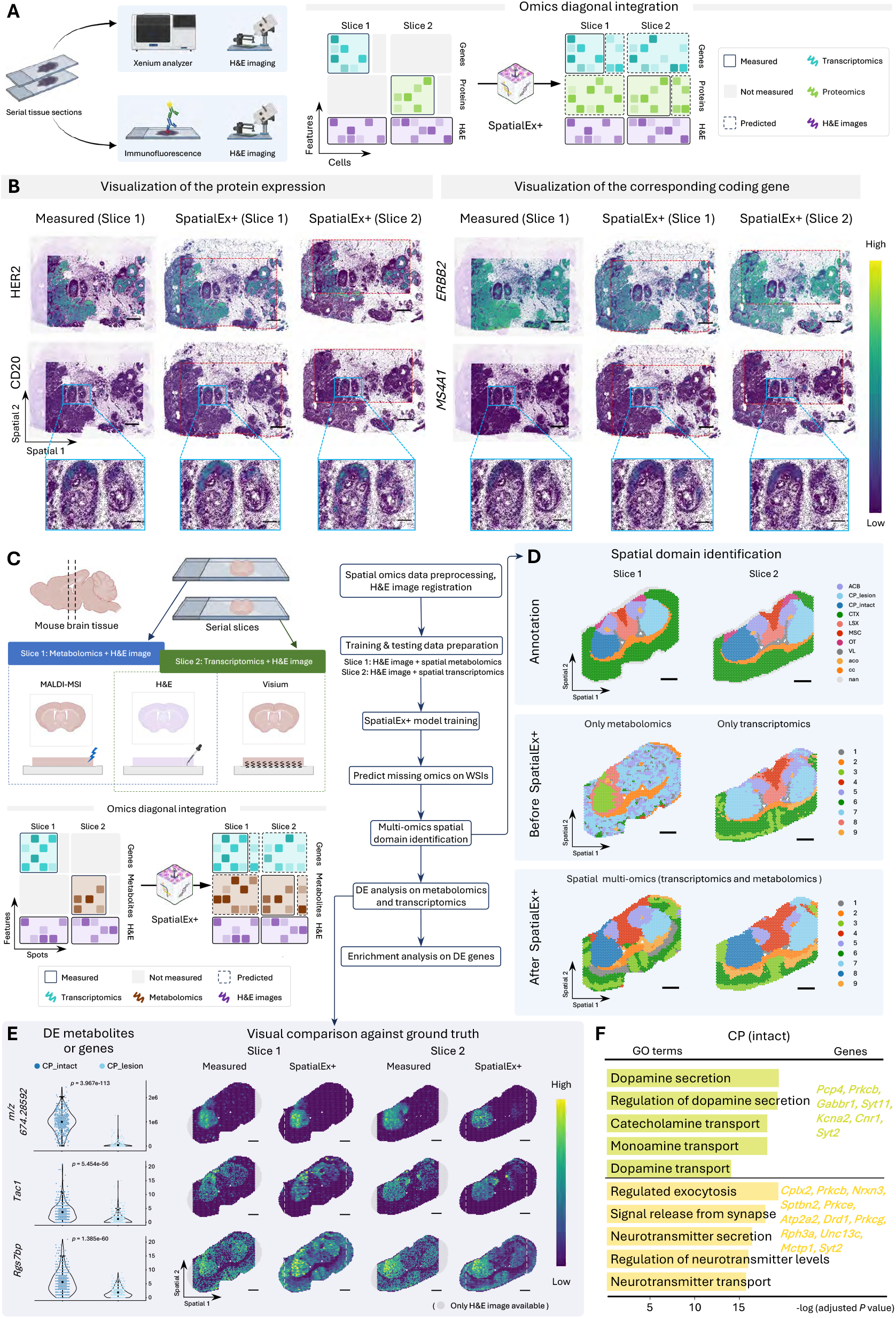
SpatialEx+ enables spatial multi-omics through omics diagonal integration. **A**. Schematic diagram for spatial transcriptomics-proteomics data diagonal integration using SpatialEx+. **B**. Visualizations of protein CD20 and HER2 and their coding genes, *MS4A1* and *ERBB2*. Measured expression on Slice 1, and whole-slide predictions of SpatialEx+ on both slices are shown. **C**. Schematic diagram of spatial transcriptomics-metabolomics data diagonal integration using SpatialEx+. **D**. Top: spatial domain annotations of both slices. Center: spatial domains identified using measured omics, metabolomics on Slice 1 and transcriptomic on Slice 2. Bottom: spatial domain identification using SpatialEx+ supplemented multi-omics for both slices. **E**. Left: scatter plots of *m/z* 674.28592 expression between lesion (***n***=478 spots) and intact (***n***=466 spots) CP regions on Slice 1, and *Tac1* and *Rgs7bp* expression lesion (***n***=420 spots) and intact (***n***=297 spots) CP regions on Slice 2. Two-sided unpaired Wilcoxon rank-sum tests were used to compare the differential, with *P* values shown. In the boxplot, the center line, center dot, box limits, and whiskers denote the median, mean, upper and lower quartiles and 1.5× interquartile range, respectively. Right: visualizations of metabolite *m/z* 674.28592, genes *Tacl* and *Rgs7bp*. **F**. GO terms enriched for the genes highly expressed within the intact CP region. The length of the bar represents the −log10 (adjusted *P* value) from one-tailed Fisher’s test. *P* values were corrected for multiple-hypothesis testing using the Benjamini–Hochberg method. (Scale bars in B: 1mm, in zoomed-in B: 250 um, in D, E: 500 um)

We then validated SpatialEx+ on the second metabolomic-transcriptomic data. The dataset consists of mouse brain slices generated using the SMA protocol^14^ (Fig. 5C). We processed the dataset to accommodate the omics diagonal integration, i.e., H&E image and metabolomics data on Slice 1, H&E image and transcriptomics data on Slice 2. Spatial metabolomics data on Slice 1 are obtained using the Matrix-Assisted Laser Desorption/Ionization Mass Spectrometry Imaging (MALDI-MSI) technique, while spatial gene expression data on Slice 2 are generated through Visium sequencing. Notably, each tissue slice comprises an intact hemisphere of healthy tissue alongside a lesioned hemisphere, modeled to represent Parkinson’s Disease (PD) through unilateral 6-hydroxydopamine (6-OHDA) administration (Fig. 5D, first row). First, we performed spatial domain identification on the metabolic data from Slice 1 and the transcriptomics data from Slice 2, respectively. The results reveal that the metabolic data is insufficient to resolve the brain anatomy structures, while using SRT data fails to distinguish the lesioned and intact choroid plexus (CP) (Fig. 5D, second row). After using SpatialEx+ on this data, each slice is equipped with both SRT and spatial metabolomics. Feeding the generated spatial multi-omics data into the state-of-the-art multi-omics analysis algorithm SpatialGLUE^38^, both the anatomic and pathological domains, including CP_lesion and CP_intact, could be identified successfully (Fig. 5D, third row), demonstrating the necessity of the generated different omics data and the utility of SpatialEx+ in a wide range analysis. A plausible reason for this phenomenon is that SpatialEx+ not only links histology and omics, but also introduces the omics cycle module to establish omics-omics associations. Quantitative performance metrics are detailed in Extended Data Fig. 8B, C.

Fig. 5E (left) quantitatively shows the expression of a selected differentially expressed (DE) metabolic *m/z* 674.28592, as well as two genes, *Tac1* and *Rgs7bp*, in CP_lesion and CP_intact. Fig. 5E (right) and Extended Data Fig. 8D, E qualitatively illustrate the spatial distribution of these DE metabolites or genes. Here, *m/z* 674.28592 has been identified as highly associated with PD by the original paper^14^, and its accurate identification is crucial for building a robust multi-omics transition model. The *Tac1* gene encodes tachykinin peptide^61^, which is believed to function as a neurotransmitter, and the *Rgs7bp* gene may be associated with risk of aspirin-exacerbated respiratory disease^62^. SpatialEx+ complements the gene and metabolite expression in the cortical areas outside the original sequencing area, enabling multi-omics data of the mouse brain across the whole section. The *m/z* 674.28592 are primarily presented in the intact caudate putamen, while the spatial expression patterns of *Tac1* and *Rgs7bp* exhibit a unilateral enrichment trend. The metabolic expressions generated by our proposed SpatialEx+ closely resemble the ground truth distribution. Specifically, the metabolic expression levels across different functional regions exhibit a high degree of similarity to the ground truth (Fig. 5E right). The predicted distributions of the two genes (*Tac1* and *Rgs7bp*) also closely align with the measured ground truth gene distributions. Moreover, we performed GO enrichment analyses on the genes in the intact region of the mouse brain sample, and the results indicated that this group of genes is associated with dopamine (Fig. 5F). The aforementioned two sets of multi-omics experiments demonstrate that by using SpatialEx+ to complete multi-omics data on the serial tissue slice, it provides a more comprehensive understanding of disease mechanisms from different biological layers.

### SpatialEx+ is robust even when there is weak or no overlap between slices

While previous sections demonstrated SpatialEx+’s capabilities on adjacent serial sections, here we evaluate its performance under more challenging conditions that violate the adjacent section requirement. Specifically, we test scenarios with significant spatial differences and sections lacking overlapping regions, conditions commonly encountered in practical applications.

To systematically evaluate robustness, we simulated different sequencing area configurations between adjacent slices using a sliding window approach, while using the actual measured data from the entire sequencing area for evaluation (Fig. 6A). Starting from the center of the maximum overlapping region between two slices (Supplementary Fig. 10A), we systematically shifted the simulated sequencing windows along opposite X-axis directions while maintaining consistent cell numbers through dynamic selection (Supplementary Fig. 10B). This generated a spectrum of test cases from fully overlapping sequencing areas (no sliding) to completely non-overlapping areas (stride 9) (Fig. 6B). Importantly, while we used these sliding windows to simulate different sequencing area configurations during training, all performance evaluations were conducted using the true measured data across the entire sequencing area.

**Fig. 6:**
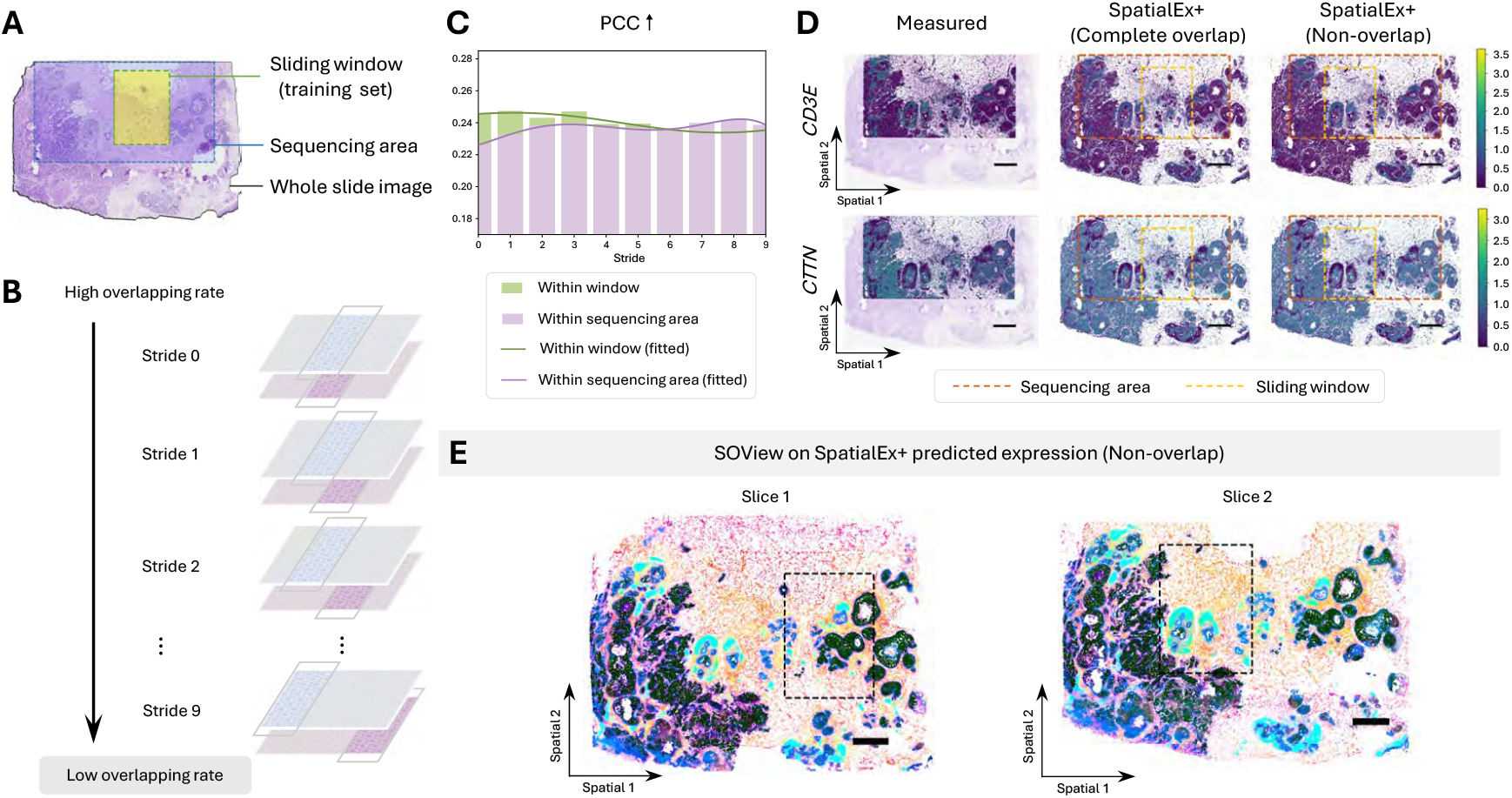
SpatialEx+ is robust even when there is weak or no overlap between slices. **A**. The definition of whole slide image, sequencing area, and sliding window. **B**. Illustration of sliding windows striding from highly to low overlapping scenarios. **C**. Bar plot of SpatialEx+’s performance under different sliding strides. Green and purple denote window-level and full-area performance, respectively. Curves represent fitted trends. **D**. Spatial visualizations of selected genes (*CD3E* and *CTTN*) at Stride 1 (complete overlap) and at Stride 9 (non-overlap) from SpatialEx+. **E**. SOView visualization of SpatialEx+ predicted expression at Stride 9 (non-overlap). (Scale bars in D, E: 1 mm)

SpatialEx+ demonstrates strong robustness, with no significant performance decline observed across varied overlap rates (evidenced by the flat trend of both within window and within sequencing area curves in Fig. 6C). This highlights the potential capability of SpatialEx+ in scenario that the sequencing area can be expanded through serial slices. The quantitative results across the entire sequencing area show only a small difference compared to the results within the window (evidenced by the small difference between purple and green bars for each stride in Fig. 6C), providing quantitative evidence of the generalizability of SpatialEx+ in extending the sequencing area to larger area. The immune cell marker *CD3E*^63^ and the tumor cell marker *CTTN*^64^ are visualized in Fig. 6D, while additional gene visualizations can be found in Extended Fig. 9. The results demonstrate that SpatialEx+ is capable of accurately reconstructing the true spatial distribution of important markers, regardless of whether the serial slices are fully overlapping or completely non-overlapping. Notably, each slice used for training contained only 20,000 to 30,000 cells within the windows, compared to over 100,000 cells across the entire sequencing area, which demonstrated that SpatialEx+ was able to generalize accurately on H&E images that were not available during training. Subsequently, under the experimental setup with no overlap (Stride 9), the gene expression predicted by SpatialEx+ on the two slices was input into Cellcharter for joint spatial domain identification. We first input the genes predicted by SpatialEx+ on the two slices into Cellcharter to generate spatially aware embeddings of each cell, and then visualize the results using SOView^65^, which is designed for visualization and analysis of high-plex spatial omics data. The spatial domain results are consistent across both slices, and the tissue structure is clearly delineated (Fig. 6E). Moreover, Supplementary Fig. 11 explores how varying sample sizes affect the performance of gene expression prediction.

## 3. Discussions

In this study, we addressed important challenges in spatial multi-omics analysis through SpatialEx and SpatialEx+, frameworks that enable high-parameter spatial profiling and multi-omics integration using histology as an anchor. Our approach overcomes key limitations of existing methods by enabling comprehensive molecular profiling without requiring complex simultaneous multi-omics co-measurements.

Our comprehensive evaluations across multiple experimental scenarios—from small-scale validations to million-cell tissue sections, from single-modality expansions to multi-omics integrations, and from ideal adjacent sections to challenging non-overlapping configurations—demonstrate SpatialEx+’s capabilities as a versatile and robust solution for spatial molecular data analysis.

However, several important challenges remain for advancing spatial diagonal integration. First, current implementations assume consistent spatial resolution across different measurements. Future developments should address cross-slice integration where spatial resolutions vary between modalities, potentially through adaptive resolution mapping, multi-scale integration approaches, or super-resolution approaches similar to iSTAR^33^.

Second, numerous methodologies have been developed for single-cell diagonal integration. A conventional approach entails projecting multimodal data into a shared feature space informed by prior knowledge (e.g., gene and coding proteins), followed by the application of single-omics analysis techniques^66–70^. More recently, some approaches have incorporated knowledge graphs into neural network frameworks to facilitate more efficient and interpretable integration^31^. Incorporating such biological prior knowledge or knowledge graphs into our framework has the potential to further improve its performance.

Third, while our framework successfully handles paired sections, extending to simultaneous integration of multiple adjacent slices (>2) would enable more comprehensive tissue analysis, requiring sophisticated multi-slice alignment strategies and expansion of the omics cycle module. Additionally, technical improvements could further enhance performance. For example, leveraging newer developments in deep hypergraph learning could better capture cellular interactions and microenvironmental information.

The growing importance of spatial multi-omics in understanding cellular behaviors and biological processes makes our framework particularly valuable for standard laboratories unable to conduct spatial multi-omics experiments directly. By lowering the barrier to spatial multi-omics analysis while maintaining high accuracy and robustness, SpatialEx+ can accelerate applications in disease modeling, drug discovery, and tissue characterization. As spatial omics technologies continue to evolve, frameworks like SpatialEx+ that enable flexible, scalable integration will become increasingly crucial for comprehensive tissue analysis.

## 4. Methods

The proposed method comprises three components—H&E foundation model, hypergraph encoder, and omics cycle module—with the first two forming SpatialEx and all three together constituting SpatialEx+. Briefly, the H&E foundation model is responsible for capturing cellular morphology features from H&E images and embedding the biologically meaningful knowledge in the feature space. The hypergraph encoder enhances cellular representations by incorporating spatial contextual information, and further refines these representations by integrating global tissue context through contrastive learning. The omics cycle module facilitates the mapping of omics-omics correlations, enabling flexible omics transition between different omics modalities. The subsequent subsections provide a detailed exposition of each module.

### H&E foundation model

Given a histology image X ∈ ℝ^*h*×*w*×3^, our objective is to encode informative morphology features of each individual cell into the feature space. To achieve this, we cropped square patches centered on each cell with a pre-defined radius *r*; the set of patches can be denoted as P ∈ *{p*_1_, *p*_2_,…, *p*_*N*_*}* with each patch *p* ∈ ℝ^*r*×*r*×3^ and *N* representing the total patch number. Subsequently, these patches were processed through a pre-trained encoder to obtain the cellular *d*-dimensional cellular representations. In this computational process, the general-purpose self-supervised pathology foundation model UNI^46^ is chosen given its exceptional feature extraction capability across a wide range pathology tasks. Notably, incorporating other advanced histology image foundation models, such as CONCH^71^, Phikon^72^, and Gigapath^73^, could also yield competitive feature representation, as demonstrated in Extended Data Fig. 10A. After the histology feature extraction, the cells are transformed from their original image space into the high-dimensional feature space, enabling efficient computation even with a mass of cells.

### Hypergraph encoder

To fully exploit spatial contextual information in terms of complex and higher-order cellular interactions, we incorporated a hypergraph structure into the computational framework. Specifically, a hypergraph can be defined as 𝒢 = {𝒱, ℰ}, where 𝒱 *= {v*_1_, *v*_2_, ⋯, *v*_*N*_*}* represents the set of nodes characterized by the extracted histology features and ℰ *= {e*_1_, *e*_2_, ⋯, *e*_*M*_*}* denotes the set of hyperedges derived from the spatial contextual relationships. Each hyperedge *ej* is a non-empty subset of nodes, *i.e*., ∅ ≠ *e*_*j*_ ⊆ 𝒱. The hypergraph structure is mathematically formulated using an incidence matrix A ∈ ℝ^*NxM*^ and a diagonal weight matrix W ∈ ℝ^*M*×*M*^. The elements in A and W indicate the node-hyperedge relationship as well as the corresponding hyperedge weights. In this study, we first identified *k* spatially neighboring cells for each cell and grouped them together to form a hyperedge. This construction process ensures each hyperedge contains multiple cells and each cell participates in multiple hyperedges. As a result, this hypergraph-based representation effectively models high-order interactions, facilitating more expressive and context-aware feature learning in the subsequent computation.

Following the hypergraph construction, we leveraged the hypergraph neural networks (HGNNs)^74,75^ for computation. Methodologically, we applied a two-stage message-passing mechanism^76,77^, which involves node-to-hyperedge (cell-to-microenvironment) and hyperedge-to-node (microenvironment-to-cell) aggregation. Specifically, in the first stage, each hyperedge representation is iteratively updated by aggregating the features of its constituent nodes. In the second stage, each node feature is refined by aggregating the updated hyperedge representations. The whole computation process can be formally expressed as follows:

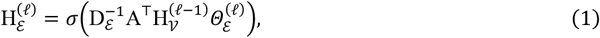

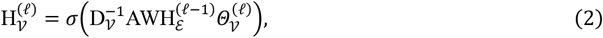

where 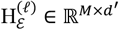 and 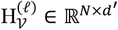 indicate the hyperedge and node representations, respectively, *σ* (·) is the Rectified Linear Unit (ReLU) function^78^, and 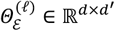 and 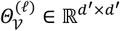 are the layer-specific trainable weights. D_ℰ_∈ ℝ^*M*×*M*^ and D_𝒱_ ∈ ℝ^*N*×*N*^ represent the hyperedge diagonal degree matrix and the node diagonal degree matrix, respectively, where 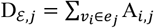 and 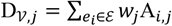.

After obtaining the encoded cellular representations, we map them to the gene expression space using a Multilayer Perceptron (MLP) decoder. Specifically, the decoder follows a two-layer linear architecture with 512 hidden units, incorporating batch normalization (BN) and the ReLU activation function, i.e., linear → ReLU → BN → linear.

Contrastive learning is known for enhancing representation learning across various classification and regression tasks^79,80^. Here, we employed contrastive learning to refine the omics-related features by encoding the global tissue context. Formally, a corrupted hypergraph is first created as 𝒢′ = {𝒱′, ℰ}, where 𝒱′ represents a set of nodes with shuffled features and ℰ is the original unchanged hyperedges. This corrupted hypergraph is then fed into the HGNN encoder, producing the corrupted representations 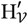. The message-passing mechanism used for this computation follows the formulation described in Equations (1) and (2). Referring the local-global contrastive framework of Deep Graph Infomax (DGI)^81^, we generated the global context representation S using an average readout operation, and maximized the mutual information between the local representation of each cell and its global context. Unlike conventional contrastive learning methods that explicitly compute loss using positive and negative sample pairs, this strategy instead associates all node embeddings—both from the original and corrupted graphs—with a single global summary vector, reducing the computational complexity from O(*N*^2^) to O(*N*). Moreover, in contrast to the standard DGI model, we opted not train a separate discriminator to assess the global information contained in the local representations and instead used a non-parametric cosine similarity objective function. In all, the global summary vector and the contrastive loss can be formulated as follows:

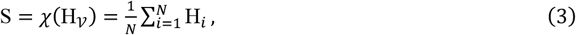

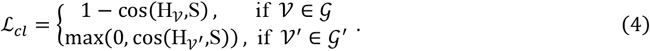

### Omics cycle module

The histology feature extractor, hypergraph representation learning, and contrastive learning jointly enable SpatialEx to translate histology to spatial omics at single-cell resolution. Building upon this foundation, SpatialEx+, an extension of SpatialEx with omics cycle modules, integrates the prediction of two distinct omics into a collaborative training framework, ensuring diagonal integration of multi-omics analysis. Specifically, each omics cycle module follows an identical structure to the MLP decoder in SpatialEx, consisting of a sequence of computations: linear → ReLU → BN → linear. It takes one type of omics data as input and translates it to another. The input omics data can either be experimentally measured or predicted from SpatialEx, enabling flexible and scalable integration. As shown in Fig. 1F, SpatialEx+ comprises omics cycle module 1 (OC_1_) and omics cycle module 2 (OC_2_). The details for module training can be found in Extended Data Fig. 1 and Methods (Model optimization). We further conducted hyperparameter sensitivity experiments on the image encoder, patch radius, and the number of spatially neighboring cells, as shown in Extended Data Fig. 10B-D.

### Model optimization

#### SpatialEx

The reconstruction loss (ℒ_*mse*_) and contrastive loss (ℒ_*cl*_) are jointly utilized to optimize SpatialEx. In terms of the first one, it targets at aligning the predicted omics expression profiles and the measured ones by minimizing the mean square errors as follows:

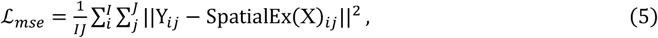

where *i* is the index for the *I* dataset-specific cells or spots and *j* is the index for the *J* genes. Y is the measured omics data. The overall loss function for SpatialEx is shown as follows:

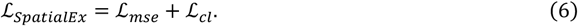

#### SpatialEx+

For optimizing SpatialEx+, we in total introduced four sets of loss functions, comprising two sets for the two SpatialEx networks and two sets for the two omics cycle modules. Here, we indicate the two SpatialEx networks as SpatialEx1 and SpatialEx2 and two omics cycle modules as OC_1_ and OC_2_. Taking the OC_1_ as an example, its optimization process involves

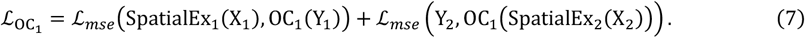

In all, the total loss for SpatialEx+ can be formulated as follows:

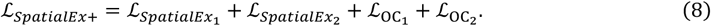

### Statistics and reproducibility

Cells and genes were filtered based on quality-control thresholds and Moran’s *I* statistics (see “Data preprocessing” section). No statistical method was used to predetermine sample size. No data were excluded from the analyses. Randomization was achieved by setting random seeds. The Investigators were not blinded to allocation during experiments and outcome assessment.

### Data preprocessing

In processing H&E images, one critical step is locating each individual cell for subsequent feature extraction. For the data labeled with cell boundaries, e.g., Xenium, we directly utilized the provided information to locate cells in images. In terms of the tissues using other techniques (not providing cell boundaries) and exceeding the sequencing area, we applied Cellpose^45^ to identify the cellular position.

#### Xenium *in situ* human and mouse data

We used two Xenium breast cancer datasets for experiments, including spatial transcriptomics-proteomics human breast cancer dataset and spatial transcriptomics human breast using the entire sample area. Both datasets comprise two consecutive Xenium slices with the panel size of 313 genes and 280 genes, respectively. Additionally, in the first dataset, Slice 1 includes a paired immunofluorescence (IF) image with markers for HER2 and CD20 proteins. Moreover, we also used Xenium datasets comprising human colon, mouse colon, and human skin tissues, each consisting of a single tissue section. The panel sizes for these datasets are 325, 379, and 282, respectively. For these datasets, we employed identical preprocessing operations for gene expressions, following the Squidpy tutorial [<u>Analyze Xenium data — squidpy documentation</u>]. Specifically, we first filtered out cells with fewer than 10 detected gene expression counts, and then used *scanpy.pp.normalize_total* to normalize the total gene expression across all cells. Afterward, we applied *scanpy.pp.log1p* for log transformation. For the IF data, we followed NicheTrans^82^ for image registration and protein quantification. This process enables the precise alignment of IF images with spatial transcriptomics data. Following this integration, each cell was characterized by a comprehensive molecular profile, incorporating both gene expression and protein expression values. Considering the small proteomics panel size, we only applied *scanpy.pp.scale* for data preprocessing. We then followed the [<u>Understanding Xenium Outputs - Official 10x Genomics Support</u>] tutorial to align the H&E image pixels to the sequenced physical locations using Xenium Explorer.

#### Spatial transcriptomics-metabolomics mouse brain data

The spatial multimodal analysis (SMA) workflow enables spatial multi-omics profiling in a single tissue section; however, the measured omics data reside on distinct coordinate systems and exhibit differing resolutions. We followed the multi-omics alignment procedures in NicheTrans to generate the fully aligned spatial multi-omics data. The input transcriptomic panel is defined as the intersection of the top 1000 spatially variable genes across the three slices, while the metabolite panel consists of the intersection of the top 50 spatially variable metabolites shared among them. We first perform log transformation for gene expression data, while leaving the metabolomic expression data unchanged. Afterwards, we scale the two types of omics data uniformly to the range of 0-1 using max-min normalization.

### Implementation details

For hypergraph construction, we set *k* as 7, identifying the 7 nearest cells along with the center cell to construct the hyperedges. Additionally, all edge weights in the hypergraph are set to 1, i.e., W is an identity matrix. We employed a two-layer (i.e., *ℓ*=2) HGNN to obtain cell representations. We trained the model using the Adam^83^ optimizer with a learning rate of 0.001, and set the number of training epochs to 500. Note that the well-trained H&E foundation model is fixed for feature extraction and will not be updated during the training process. All experiments were conducted on one 24 GB Nvidia GeForce RTX 3090, using PyTorch 2.3.1 and Python 3.8 environment.

#### SOView visualization

We conducted joint spatial domain identification on two slices using the aligned gene expression using Cellcharter. Specifically, the process begins by applying scVI^84^ for dimensionality reduction to obtain a low-dimensional representation. Next, it aggregates representations from varying numbers of neighboring cells based on their spatial locations, and concatenates them to create spatially-aware embeddings for each cell. These embeddings are then input into a Gaussian mixture model for clustering. To visualize the spatially-aware embeddings, we employed SOView^65^. The first three components of the UMAP embeddings are rescaled to the range 0–255, forming an RGB color matrix. A color plot is then generated based on this RGB color matrix and the spatial location matrix. In this way, the color similarities between spots reflect the similarities in gene expression, allowing tissue heterogeneity to be visualized in a single plot.

#### Cell type transferring

We jointly performed Principal Component Analysis (PCA) on the measured gene expressions of Slice 1 and the predicted gene expression of Slice 2, extracting 50 Principal Components (PCs). Afterwards, the data of Slice 1 was utilized to optimize the MLP classifier, with the PCs serving as inputs and cell type annotations as targets. Here, the MLP classifier is constructed by three linear layers, where the first two layers are followed by a LeakyReLU activation function and the last layer followed by a Softmax function. Considering the inherent long-tail distribution of different cell types, focal loss^85^ was utilized to optimize the MLP classifier. Additionally, cell types with limited samples were augmented by adding Gaussian noise, maintaining balanced class distribution in each batch. The model parameters were updated using the Adam optimizer with a learning rate of 0.001 and no weight decay.

To thoroughly evaluate transferability, we performed 10-fold cross-validation on Slice 1. In each fold, nine parts were used for training and the remaining part for validation. The best-performing model on the validation set was then tested on Slice 2.

#### GO enrichment analysis

We first used the *scanpy.tl.rank_genes_groups*() function from Scanpy to rank gene importance within each spatial domain. Genes with a logfoldchange > 1.5 were selected as marker genes for each spatial domain. These marker genes were then converted to gene IDs and subjected to GO functional enrichment using the *enrichGO* function from the R package clusterProfiler. Multiple testing correction was performed using the Benjamini-Hochberg FDR method (pAdjustMethod = ‘fdr’), with *P* value and *Q* value thresholds of 0.05 and 0.2, respectively.

### Evaluation

In this study, three evaluation metrics, including Pearson correlation coefficients (PCC), correlation matrix distance (CMD), and the structural similarity index measure (SSIM), Moran’s *I* and Spearman’s rank correlation coefficient (SPCC) were utilized for quantitatively assessing the model’s performance.

**PCC** serves to quantify the linear correlation between two variables, and its definition is shown as follows:

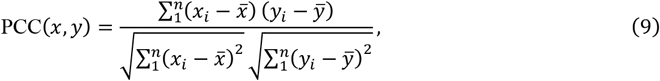

where *xi* represents the expression of gene *i*. 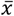 and 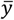 denotes the mean values of *x* and *y*.

**CMD** is a general measurement of the difference between two correlation matrices R1 and R2. A lower CMD value indicates better results. CMD can be formulated as follows:

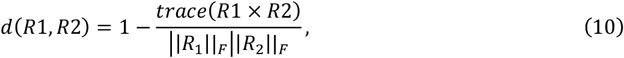

where *trace*(.) calculates the trace of the matrix, and || · ||_*F*_ represents the Frobenius norm of the matrix.

**SSIM** is used to evaluate the similarity between the ground truth and the predicted spatial structure of gene expression, with higher values indicating greater similarity. In this study, we used SSIM to evaluate the structural similarity of hypergraph formed by spatial information of cells. Specifically, it can be formulated as follows:

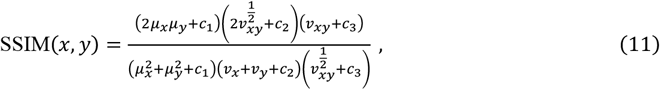

where *x* and *y* denote measured and predicted gene expression, respectively. The *μ*_*x*_ and *μ*_*y*_ represent the spatially Gaussian-filtered gene expression of *x* and *y*. Correspondingly, *vx* and *vy* indicate the spatially Gaussian-filtered variance of *x* and *y*, while *vxy* is the spatially Gaussian-filtered covariance between *x* and *y*. The *c1*, and *c2* are predefined constant, *c*_1_ *=* (0.01 (*max* (*x*) *- min* (*x*))) ^2^, *c*_2_ *=* (0.03 (*max* (*x*) *- min* (*x*))) ^2^ and *c =* (0.03 (*max* (*x*) *- min* (*x*))) ^2^ and 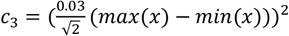.

**Moran’s *I*** is a measure of spatial autocorrelation, reflecting the degree to which similar values cluster in space. It is defined as:

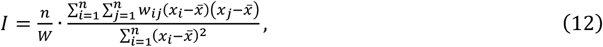

where *x*_*i*_is the expression level at location *i*, 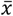 is the mean expression across all locations, *w*_*ij*_ is the spatial weight between locations *i* and *j, n* is the number of spatial units, and *W =* Σ_*i*,*j*_ *w*_*ij*_ is the sum of all spatial weights.

**SPCC** is used to assess the monotonic relationship between two variables. In our study, SPCC is calculated between the Moran’s *I* values of predicted and measured gene expression to evaluate how well spatial autocorrelation is preserved. It is defined as:

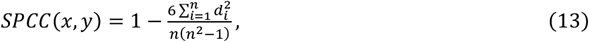

where 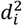 is the difference between the ranks of gene *i* in the predicted and measured Moran’s *I* lists, and *n* is the total number of genes.

## Data availability

The datasets used in this study are publicly accessible, with detailed statistical information summarized in Supplementary Table 1. The Xenium Human Breast Cancer tissue dataset is available at https://www.10xgenomics.com/products/xenium-in-situ/human-breast-dataset-explorer. The Xenium Human Colon tissue dataset is available at https://www.10xgenomics.com/datasets/human-colon-preview-data-xenium-human-colon-gene-expression-panel-1-standard. The Xenium Mouse Colon tissue dataset is available at https://www.10xgenomics.com/cn/datasets/fresh-frozen-mouse-colon-with-xenium-multimodal-cell-segmentation-1-standard. The Xenium Human Skin tissue is available at https://www.10xgenomics.com/cn/datasets/human-skin-preview-data-xenium-human-skin-gene-expression-panel-1-standard. The Xenium Human Breast Using the Entire Sample Area dataset is publicly available at https://www.10xgenomics.com/datasets/ffpe-human-breast-using-the-entire-sample-area-1-standard. The Visium Spatial Multimodal Analysis dataset is available at https://data.mendeley.com/datasets/w7nw4km7xd/1. Source data for this study are available with this paper, and in the Zenodo repository at https://doi.org/10.5281/zenodo.17191222.

## Code availability

Our code is available on GitHub at https://github.com/KEAML-JLU/SpatialEx. The implementations of Spatial and SpatialEx+ as well as tutorials can be found at https://spatialex-tutorials.readthedocs.io/en/latest.

## Author contributions

Z.Y., X.F., and R.G. conceived and supervised the study, secured funding, and provided analytical guidance. Y.L. implemented the algorithm with assistance from C.W. and Z.W. C.W. performed the main analyses with support from Y.L. and Z.W. C.W. generated the figures with input from Z.W. and L.C. Z.L. conducted the tissue annotation. J.S., B.-Z.Q., Q.Z., and R.G. provided valuable suggestions on model design. Z.Y., X.F., R.G., Y.L., and Z.W. wrote the manuscript with the input from all authors. All authors reviewed and approved the final version of the manuscript.

## Acknowledgements

Our work is supported by National Key R&D Program of China (No. 2023YFF1204800 (Z.Y.)), National Nature Science Foundation of China (No. 32470706 (Z.Y.), No. 62303119 (Z.Y), No. 62172187 (R.G.) and No.62372209 (X.F.)), the Computational Biology Program (25JS2850200 (Z.Y.)) of Science and Technology Commission of Shanghai Municipality (STCSM), Chenguang Program of Shanghai Education Development Foundation and Shanghai Municipal Education Commission (No. 22CGA02 (Z.Y.)), Shanghai Municipal Science and Technology Major Project (No. 2023SHZDZX02 (B.-Z.Q.)), and Fund of Fudan University and Cao’ejiang Basic Research (No. 24FCA10 (Z.Y.)).

---

**Extended Figure 1.**
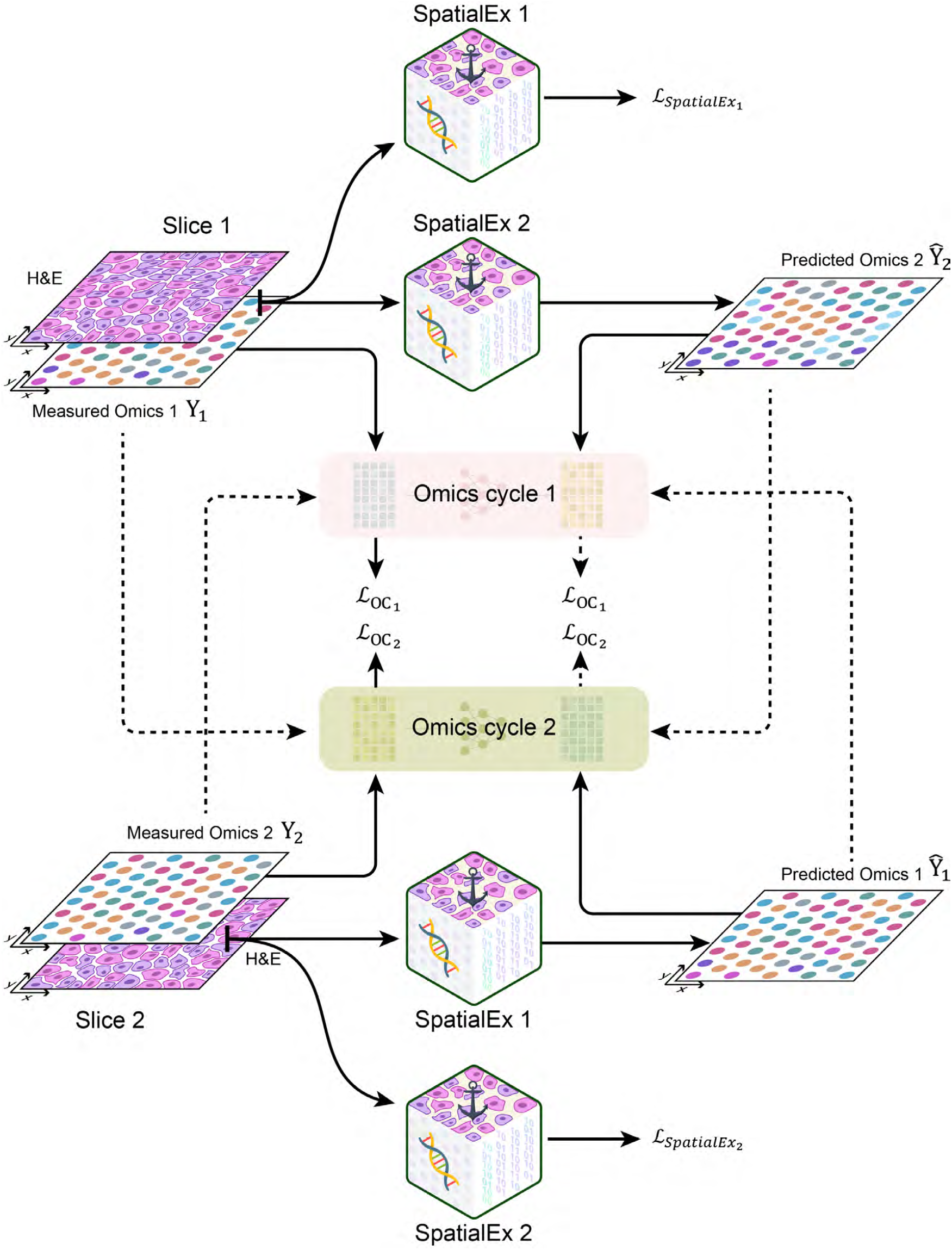

**Extended Figure 2.**
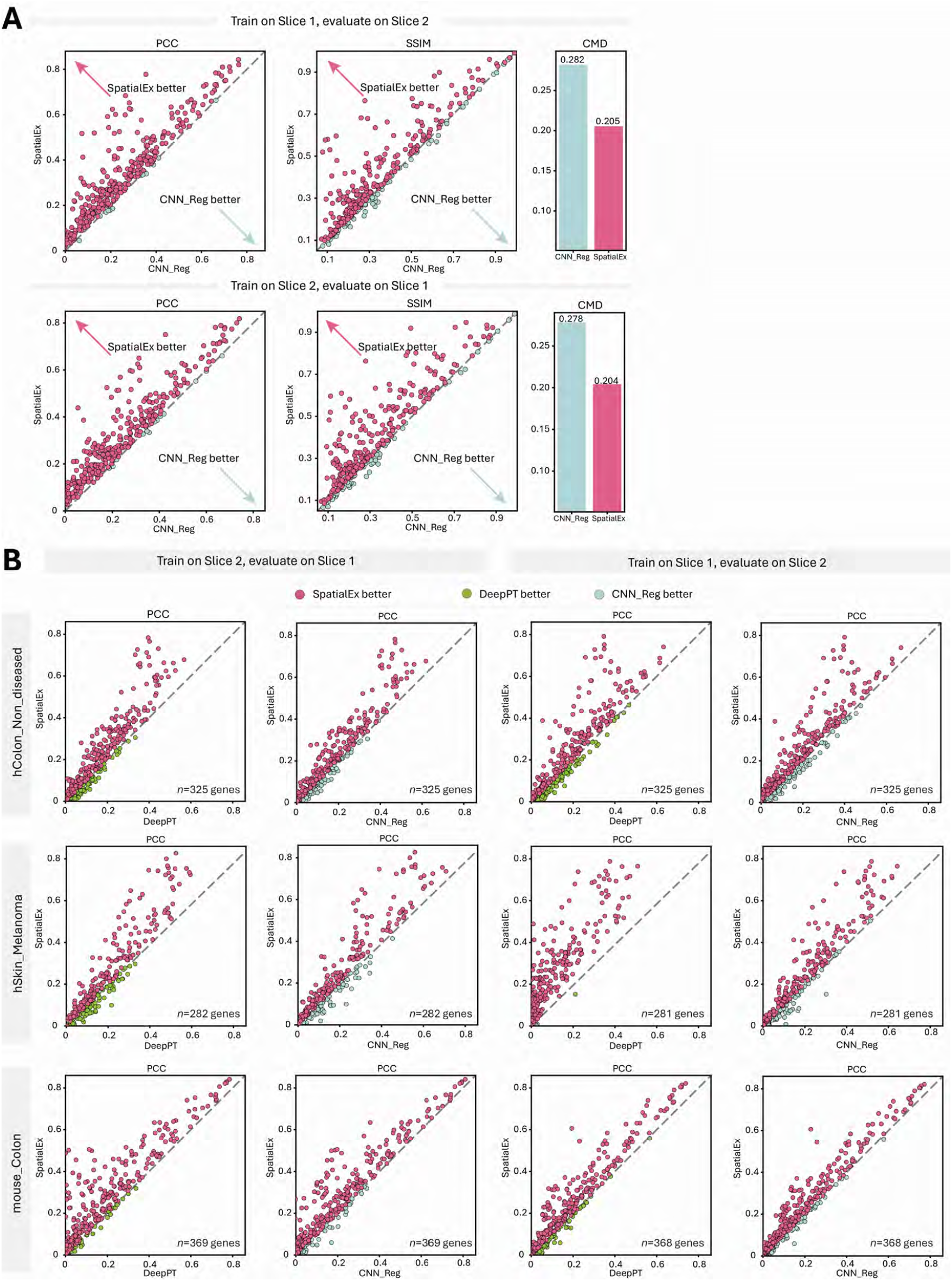

**Extended Figure 3.**
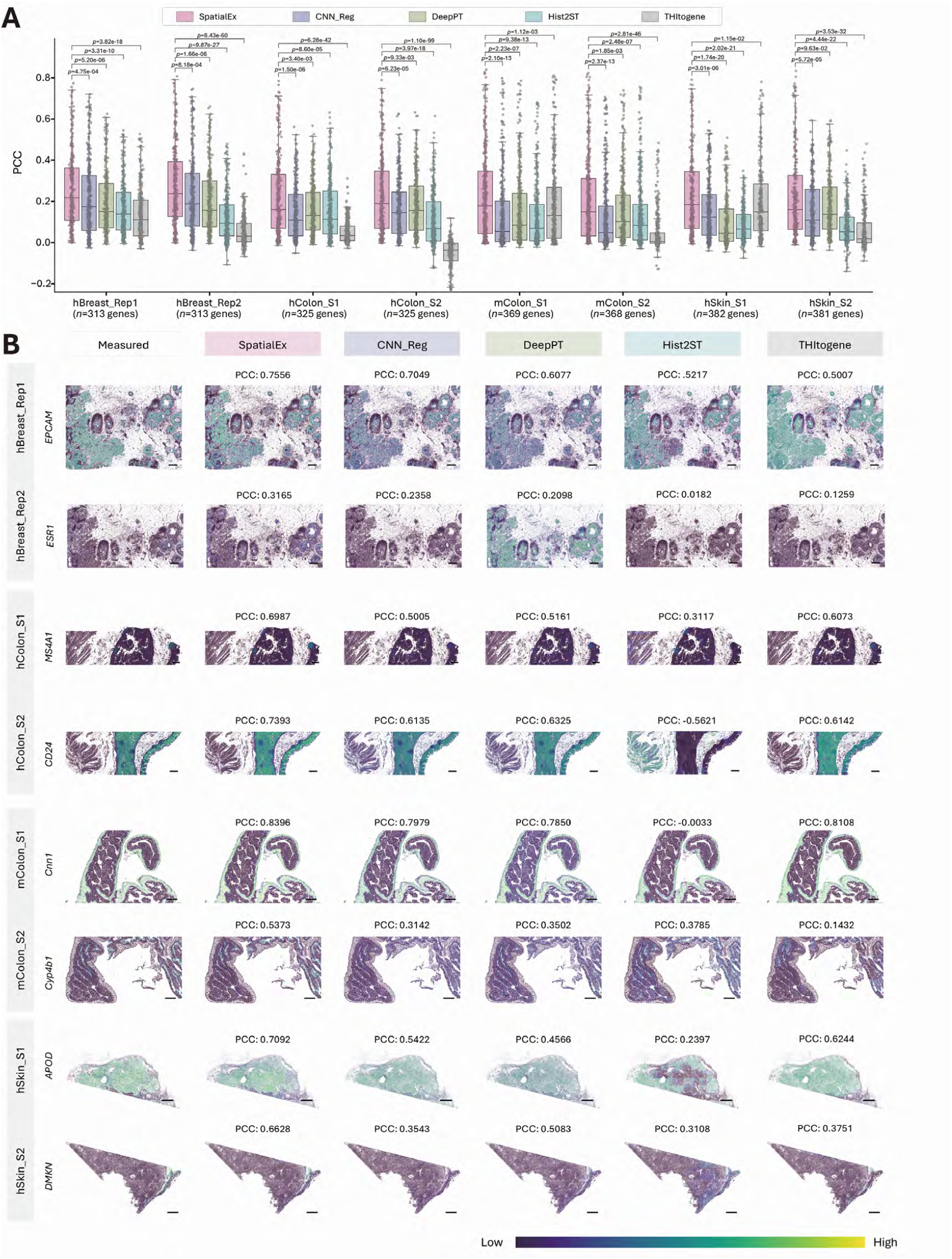

**Extended Figure 4.**
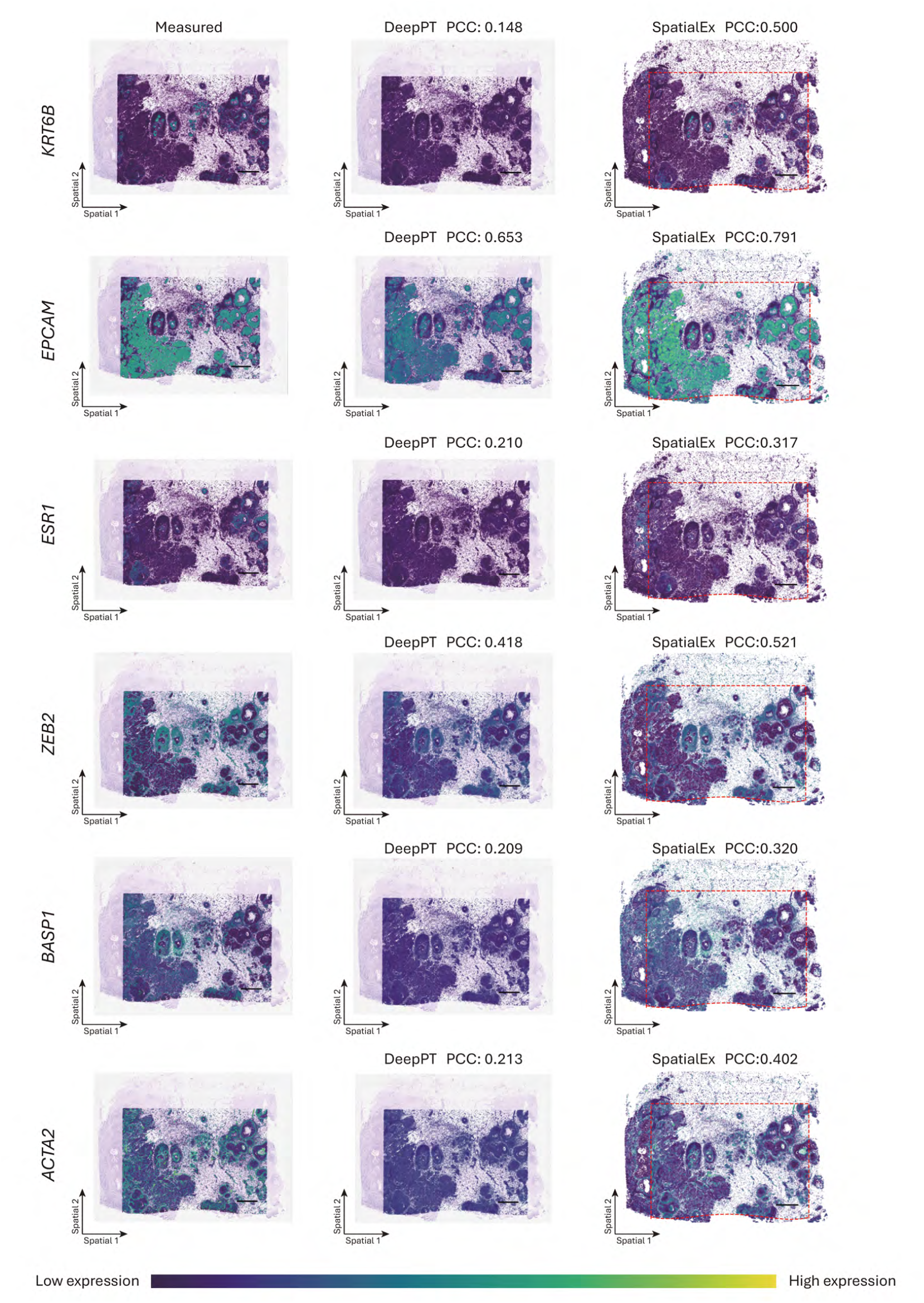

**Extended Figure 5.**
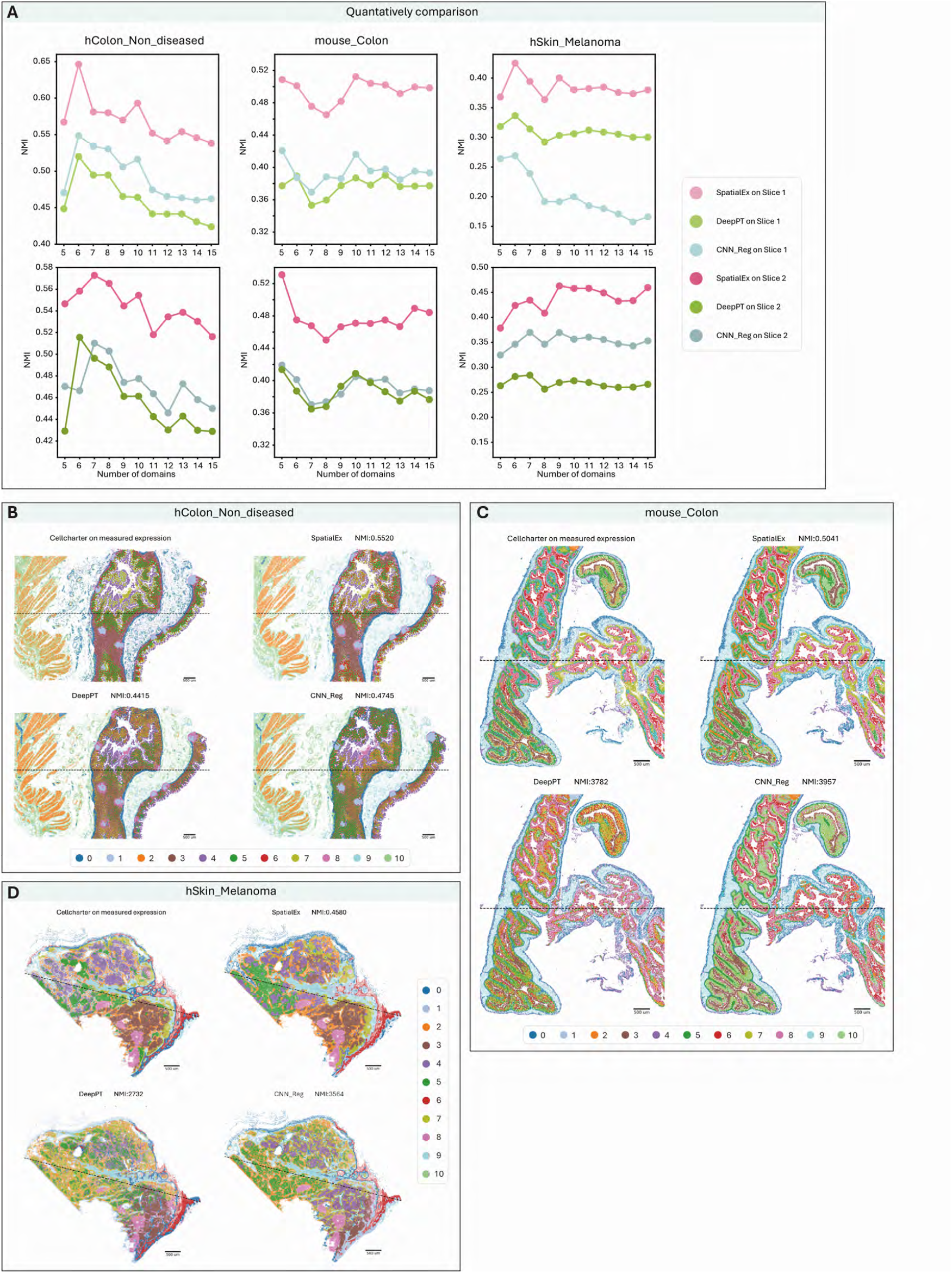

**Extended Figure 6.**
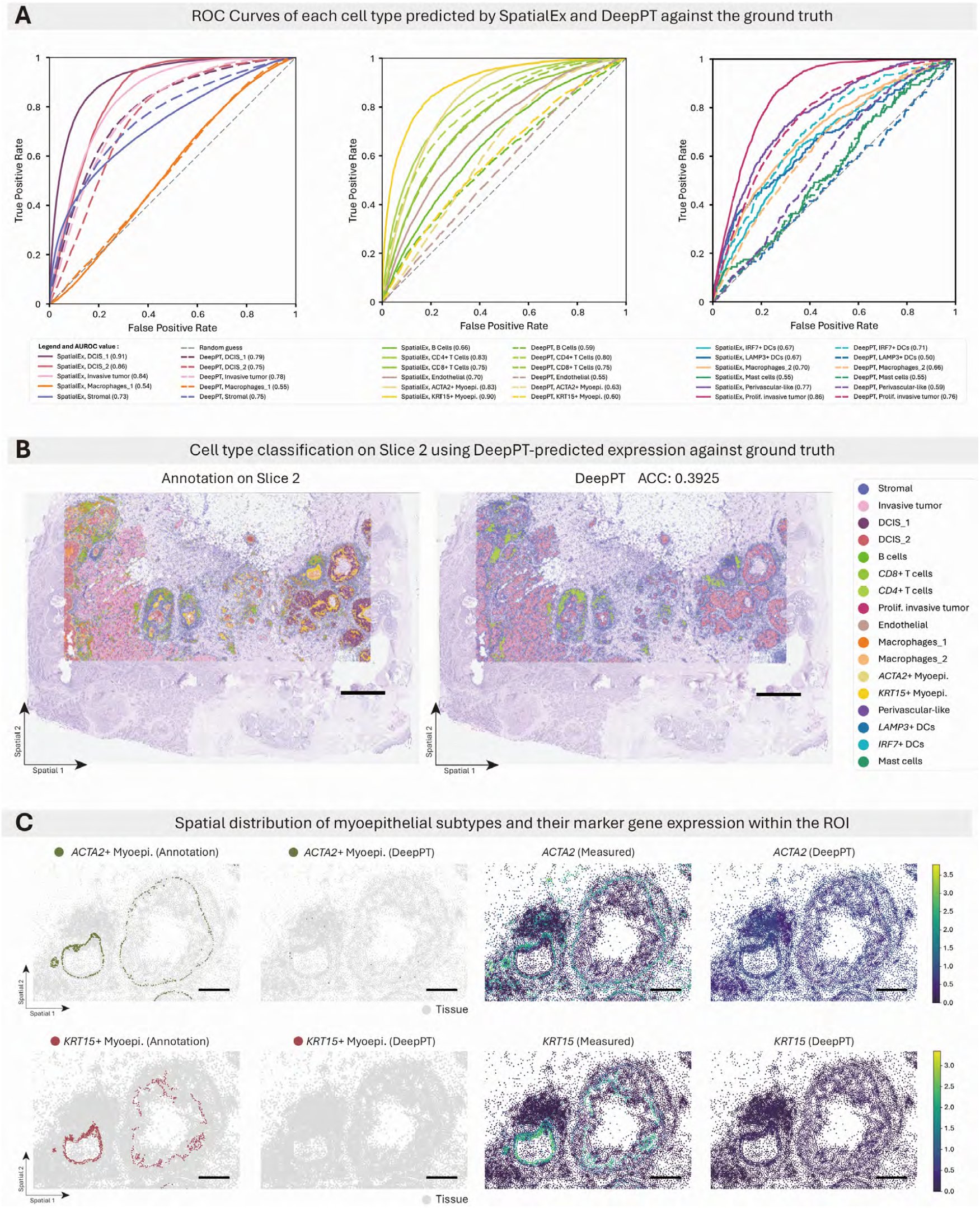

**Extended Figure 7.**
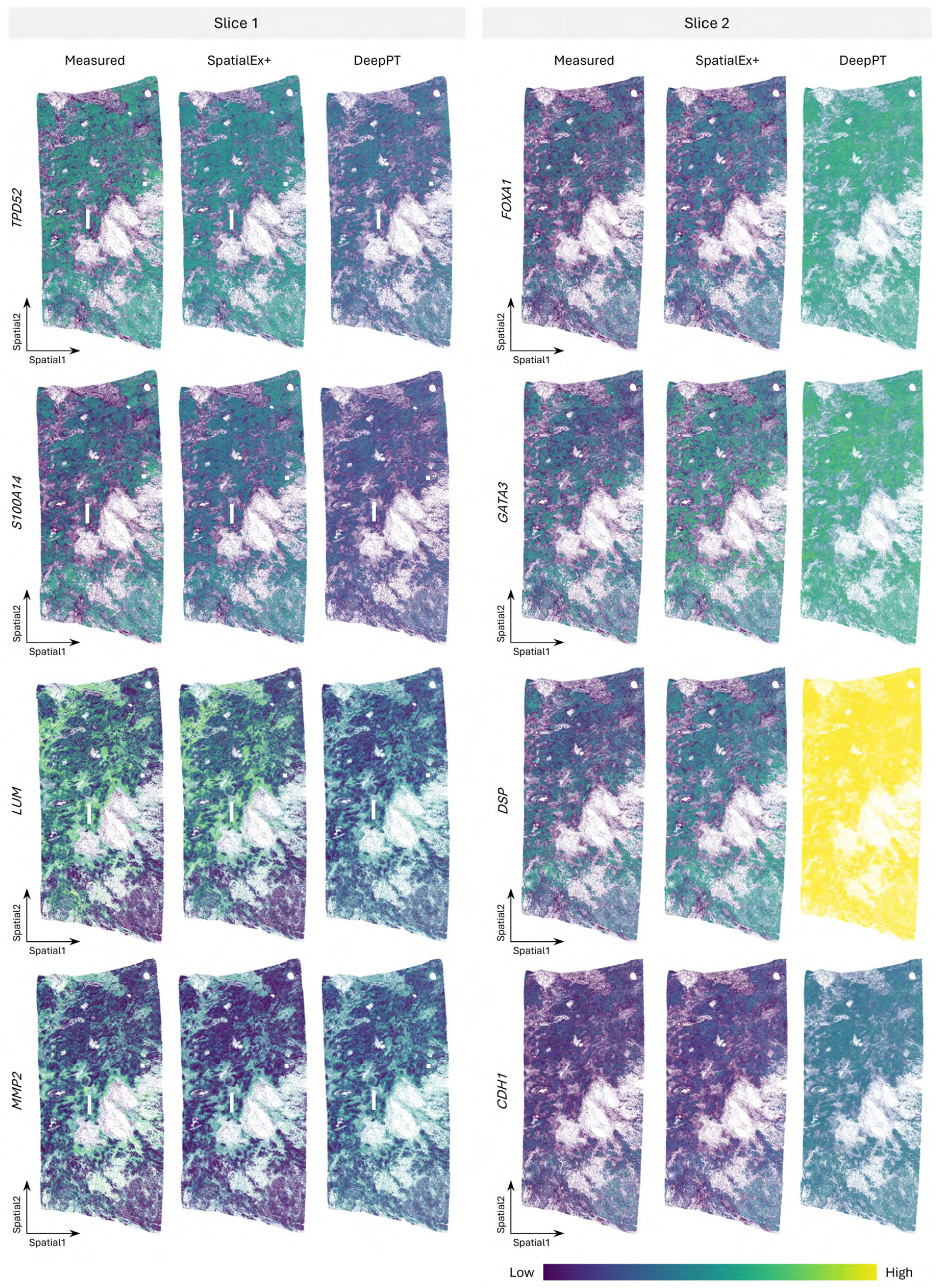

**Extended Figure 8.**
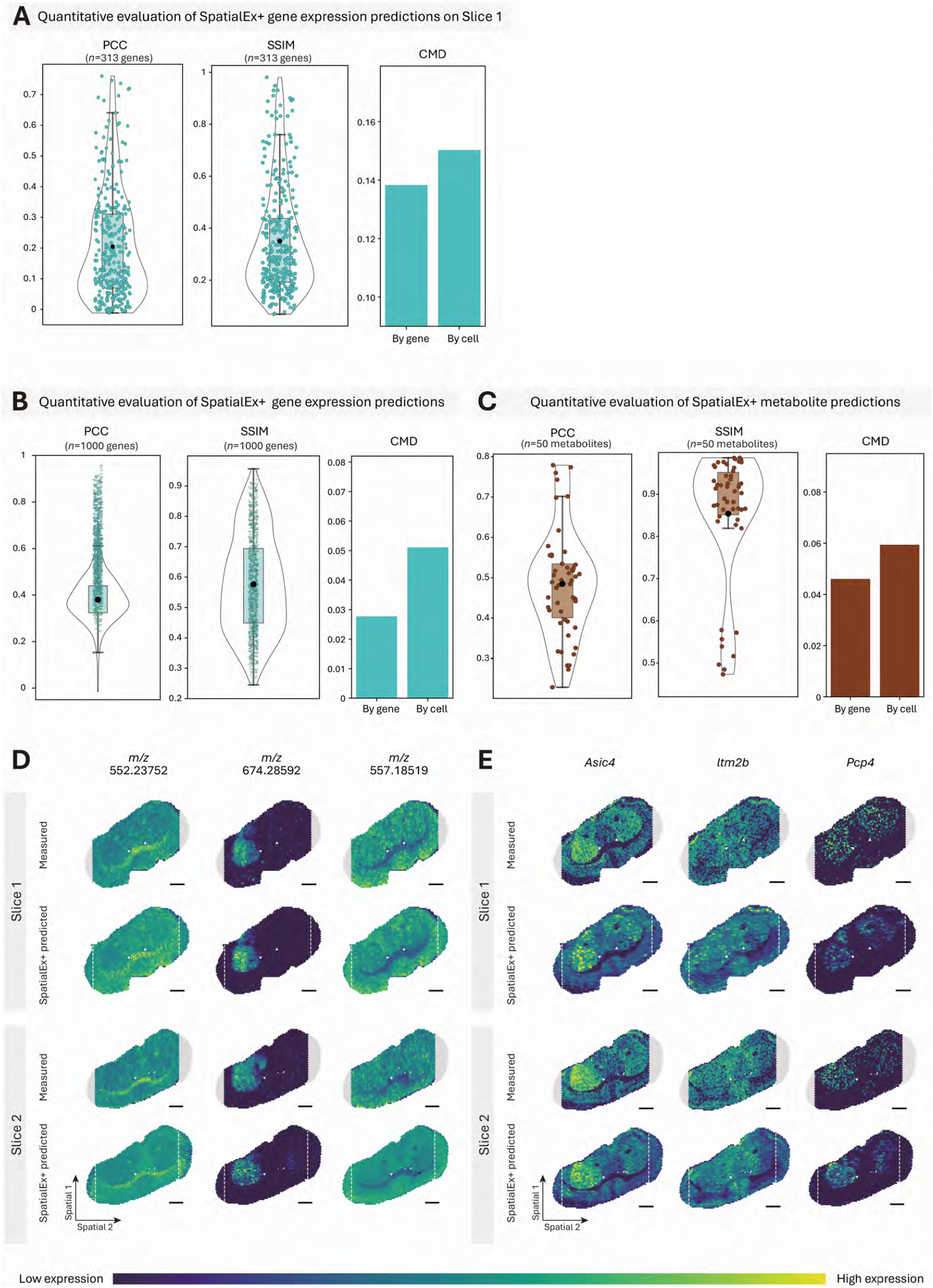

**Extended Figure 9.**
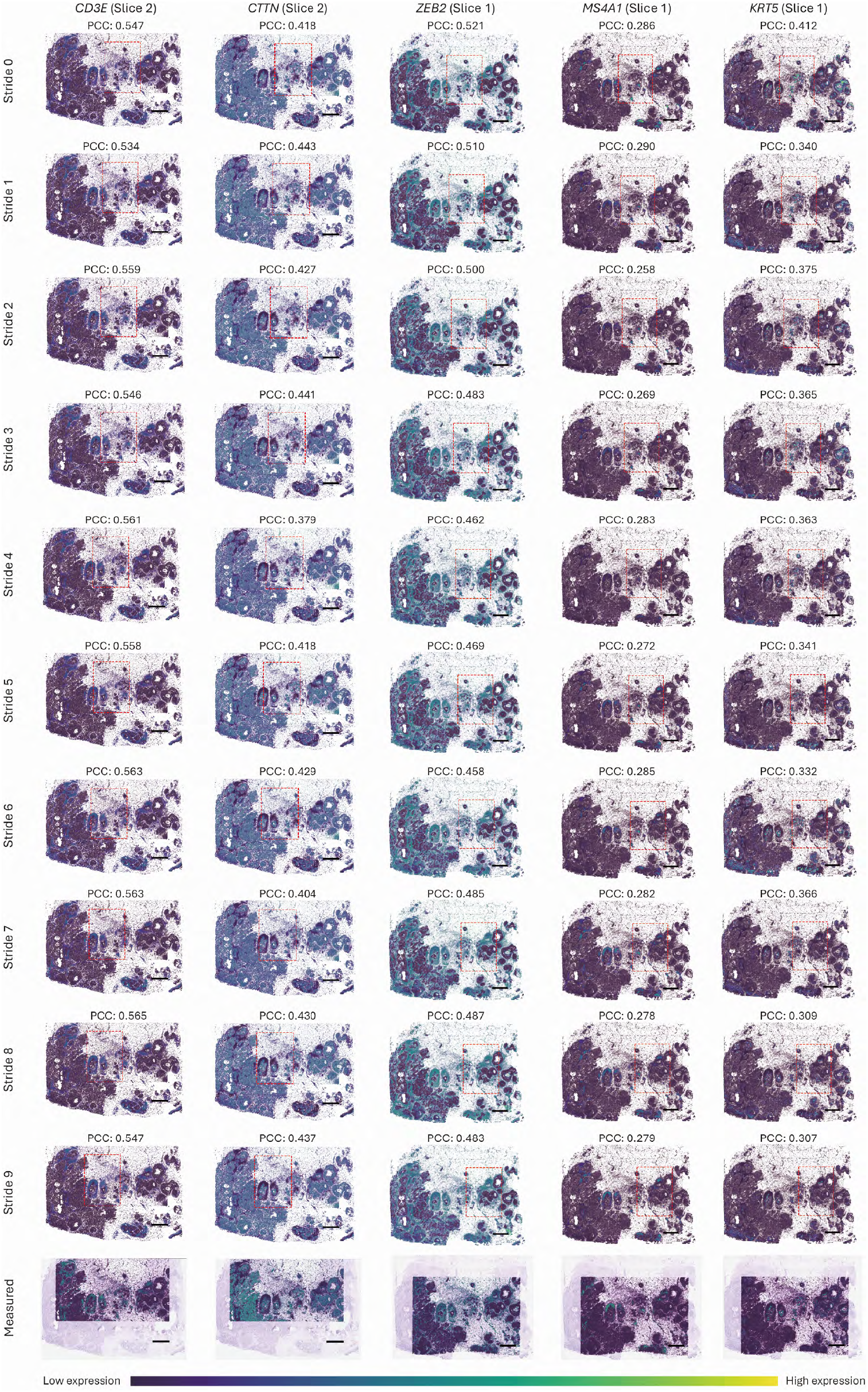

**Extended Figure 10.**
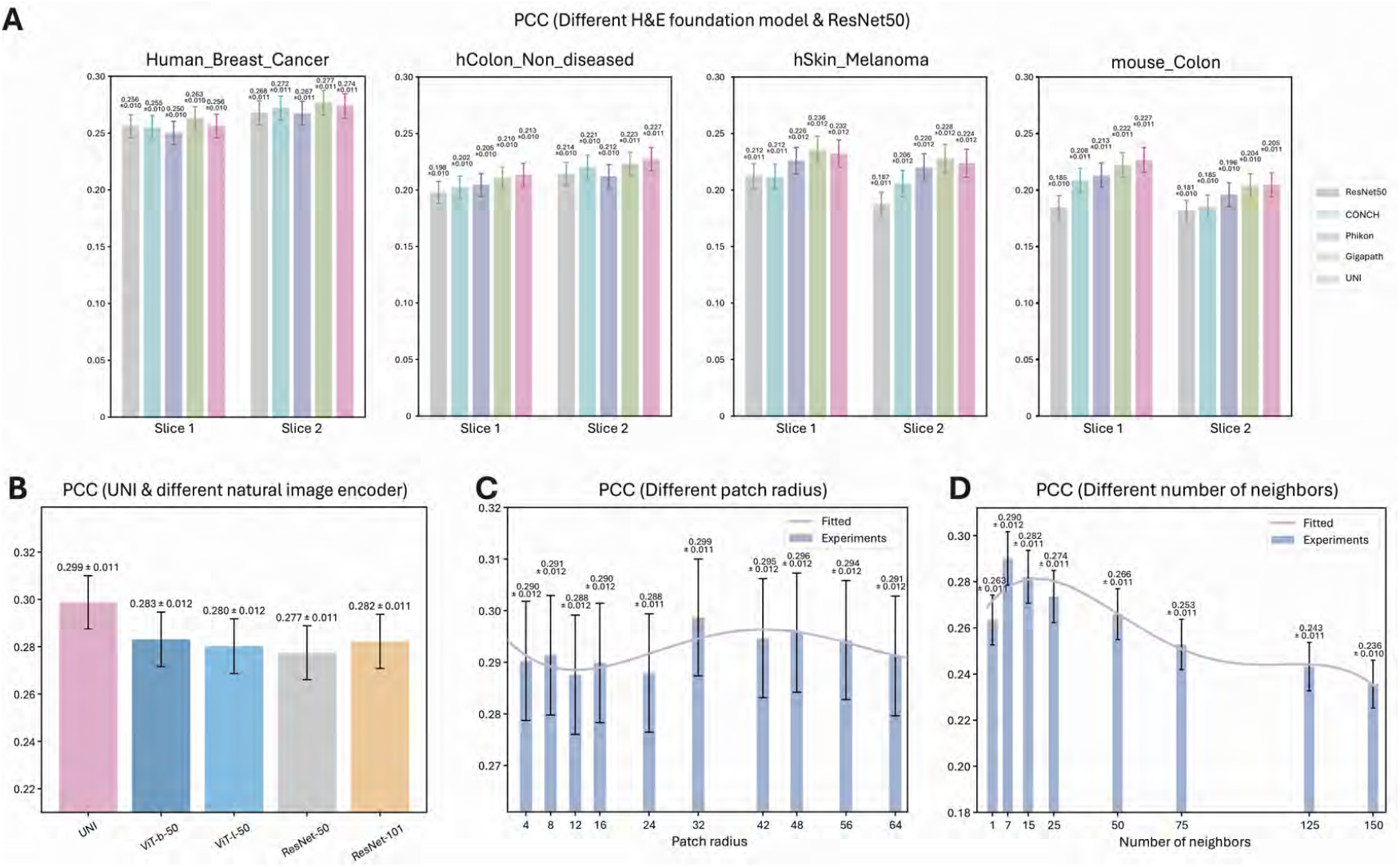

## References

1. Rao, A., Barkley, D., França, G. S. & Yanai, I. Exploring tissue architecture using spatial transcriptomics. Nature 596, 211–220 (2021).

2. Moffitt, J. R., Lundberg, E. & Heyn, H. The emerging landscape of spatial profiling technologies. Nat. Rev. Genet. 23, 741–759 (2022).

3. Mund, A., Brunner, A.-D. & Mann, M. Unbiased spatial proteomics with single-cell resolution in tissues. Mol. Cell 82, 2335–2349 (2022).

4. Bressan, D., Battistoni, G. & Hannon, G. J. The dawn of spatial omics. Science 381, eabq4964 (2023).

5. Yayon, N. et al. A spatial human thymus cell atlas mapped to a continuous tissue axis. Nature 635, 708–718 (2024).

6. Xu, Y. et al. A single-cell transcriptome atlas profiles early organogenesis in human embryos. Nat. Cell Biol. 25, 604–615 (2023).

7. Mo, C.-K. et al. Tumour evolution and microenvironment interactions in 2D and 3D space. Nature 634, 1178–1186 (2024).

8. Lucas, C.-H. G. et al. Spatial genomic, biochemical and cellular mechanisms underlying meningioma heterogeneity and evolution. Nat. Genet. 1–13 (2024).

9. Greenwald, A. C. et al. Integrative spatial analysis reveals a multi-layered organization of glioblastoma. Cell 187, 2485-2501. e26 (2024).

10. Chen, X. et al. Whole-cortex in situ sequencing reveals input-dependent area identity. Nature 1–10 (2024).

11. Klughammer, J. et al. A multi-modal single-cell and spatial expression map of metastatic breast cancer biopsies across clinicopathological features. Nat. Med. 30, 3236–3249 (2024).

12. Liu, Y. et al. High-spatial-resolution multi-omics sequencing via deterministic barcoding in tissue. Cell 183, 1665–1681 (2020).

13. Liu, Y. et al. High-plex protein and whole transcriptome co-mapping at cellular resolution with spatial CITE-seq. Nat. Biotechnol. 41, 1405–1409 (2023).

14. Vicari, M. et al. Spatial multimodal analysis of transcriptomes and metabolomes in tissues. Nat. Biotechnol. 42, 1046–1050 (2024).

15. Zhang, D. et al. Spatial epigenome–transcriptome co-profiling of mammalian tissues. Nature 616, 113–122 (2023).

16. Guo, P. et al. Multiplexed spatial mapping of chromatin features, transcriptome and proteins in tissues. Nat. Methods 1–10 (2025).

17. Vandereyken, K., Sifrim, A., Thienpont, B. & Voet, T. Methods and applications for single-cell and spatial multi-omics. Nat. Rev. Genet. 24, 494–515 (2023).

18. Deng, Y., Bai, Z. & Fan, R. Microtechnologies for single-cell and spatial multi-omics. Nat. Rev. Bioeng. 1, 769–784 (2023).

19. Tang, Z. et al. Search and match across spatial omics samples at single-cell resolution. Nat. Methods 1–12 (2024).

20. Ekvall, M. et al. Spatial landmark detection and tissue registration with deep learning. Nat. Methods 21, 673–679 (2024).

21. Wahle, P. et al. Multimodal spatiotemporal phenotyping of human retinal organoid development. Nat. Biotechnol. 41, 1765–1775 (2023).

22. Carstens, J. L. et al. Spatial multiplexing and omics. Nat. Rev. Methods Primer 4, 54 (2024).

23. Zormpas, E., Queen, R., Comber, A. & Cockell, S. J. Mapping the transcriptome: Realizing the full potential of spatial data analysis. Cell 186, 5677–5689 (2023).

24. Chen, K. H., Boettiger, A. N., Moffitt, J. R., Wang, S. & Zhuang, X. Spatially resolved, highly multiplexed RNA profiling in single cells. Science 348, aaa6090 (2015).

25. Janesick, A. et al. High resolution mapping of the tumor microenvironment using integrated single-cell, spatial and in situ analysis. Nat. Commun. 14, 8353 (2023).

26. Bonev, B. et al. Opportunities and challenges of single-cell and spatially resolved genomics methods for neuroscience discovery. Nat. Neurosci. 27, 2292–2309 (2024).

27. Piwecka, M., Rajewsky, N. & Rybak-Wolf, A. Single-cell and spatial transcriptomics: deciphering brain complexity in health and disease. Nat. Rev. Neurol. 19, 346–362 (2023).

28. Seferbekova, Z., Lomakin, A., Yates, L. R. & Gerstung, M. Spatial biology of cancer evolution. Nat. Rev. Genet. 24, 295–313 (2023).

29. Li, B. et al. Benchmarking spatial and single-cell transcriptomics integration methods for transcript distribution prediction and cell type deconvolution. Nat. Methods 19, 662–670 (2022).

30. Argelaguet, R., Cuomo, A. S., Stegle, O. & Marioni, J. C. Computational principles and challenges in single-cell data integration. Nat. Biotechnol. 39, 1202–1215 (2021).

31. Cao, Z.-J. & Gao, G. Multi-omics single-cell data integration and regulatory inference with graph-linked embedding. Nat. Biotechnol. 40, 1458–1466 (2022).

32. He, B. et al. Integrating spatial gene expression and breast tumour morphology via deep learning. Nat. Biomed. Eng. 4, 827–834 (2020).

33. Zhang, D. et al. Inferring super-resolution tissue architecture by integrating spatial transcriptomics with histology. Nat. Biotechnol. 1–6 (2024).

34. Wang, C. et al. Benchmarking the translational potential of spatial gene expression prediction from histology. Nat. Commun. 16, 1544 (2025).

35. Xu, Y. & McCord, R. P. Diagonal integration of multimodal single-cell data: potential pitfalls and paths forward. Nat. Commun. 13, 3505 (2022).

36. Chen, S. et al. Integration of spatial and single-cell data across modalities with weakly linked features. Nat. Biotechnol. 42, 1096–1106 (2024).

37. Samaran, J., Peyré, G. & Cantini, L. scConfluence: single-cell diagonal integration with regularized Inverse Optimal Transport on weakly connected features. Nat. Commun. 15, 7762 (2024).

38. Long, Y. et al. Deciphering spatial domains from spatial multi-omics with SpatialGlue. Nat. Methods 1–10 (2024).

39. Coleman, K. et al. Resolving tissue complexity by multimodal spatial omics modeling with MISO. Nat. Methods 1–9 (2025).

40. Varrone, M., Tavernari, D., Santamaria-Martínez, A., Walsh, L. A. & Ciriello, G. CellCharter reveals spatial cell niches associated with tissue remodeling and cell plasticity. Nat. Genet. 56, 74–84 (2024).

41. Singhal, V. et al. BANKSY unifies cell typing and tissue domain segmentation for scalable spatial omics data analysis. Nat. Genet. 56, 431–441 (2024).

42. Yan, X. et al. Mosaic integration of spatial multi-omics with SpaMosaic. bioRxiv 2024–10 (2024).

43. Coleman, K., Schroeder, A. & Li, M. Unlocking the power of spatial omics with AI. Nat. Methods 21, 1378–1381 (2024).

44. Chen, X. et al. HE2Gene: image-to-RNA translation via multi-task learning for spatial transcriptomics data. Bioinformatics 40, (2024).

45. Stringer, C., Wang, T., Michaelos, M. & Pachitariu, M. Cellpose: a generalist algorithm for cellular segmentation. Nat. Methods 18, 100–106 (2021).

46. Chen, R. J. et al. Towards a general-purpose foundation model for computational pathology. Nat. Med. 30, 850–862 (2024).

47. Hu, Y. et al. Unsupervised and supervised discovery of tissue cellular neighborhoods from cell phenotypes. Nat. Methods 21, 267–278 (2024).

48. Ren, H., Walker, B. L., Cang, Z. & Nie, Q. Identifying multicellular spatiotemporal organization of cells with SpaceFlow. Nat. Commun. 13, 4076 (2022).

49. Dong, K. & Zhang, S. Deciphering spatial domains from spatially resolved transcriptomics with an adaptive graph attention auto-encoder. Nat. Commun. 13, 1739 (2022).

50. Moses, L. & Pachter, L. Museum of spatial transcriptomics. Nat. Methods 19, 534–546 (2022).

51. Wilk, A. J., Shalek, A. K., Holmes, S. & Blish, C. A. Comparative analysis of cell–cell communication at single-cell resolution. Nat. Biotechnol. 42, 470–483 (2024).

52. Palla, G., Fischer, D. S., Regev, A. & Theis, F. J. Spatial components of molecular tissue biology. Nat. Biotechnol. 40, 308–318 (2022).

53. Zeng, Y. et al. Spatial transcriptomics prediction from histology jointly through transformer and graph neural networks. Brief. Bioinform. 23, bbac297 (2022).

54. Xie, R. et al. Spatially Resolved Gene Expression Prediction from Histology Images via Bi-modal Contrastive Learning. Adv. Neural Inf. Process. Syst. 36, (2024).

55. Hoang, D.-T. et al. A deep-learning framework to predict cancer treatment response from histopathology images through imputed transcriptomics. Nat. Cancer 5, 1305–1317 (2024).

56. Jia, Y., Liu, J., Chen, L., Zhao, T. & Wang, Y. THItoGene: a deep learning method for predicting spatial transcriptomics from histological images. Brief. Bioinform. 25, bbad464 (2024).

57. Qing, J. et al. Suppression of anchorage-independent growth and matrigel invasion and delayed tumor formation by elevated expression of fibulin-1D in human fibrosarcoma-derived cell lines. Oncogene 15, 2159–2168 (1997).

58. Twal, W. O. et al. Fibulin-1 suppression of fibronectin-regulated cell adhesion and motility. J. Cell Sci. 114, 4587–4598 (2001).

59. Pupa, S. M. et al. Regulation of breast cancer response to chemotherapy by fibulin-1. Cancer Res. 67, 4271–4277 (2007).

60. Kelemen, L. E. et al. Genetic variation in stromal proteins decorin and lumican with breast cancer: investigations in two case-control studies. Breast Cancer Res. 10, 1–11 (2008).

61. Yu, B. et al. TAC1, a major quantitative trait locus controlling tiller angle in rice. Plant J. 52, 891–898 (2007).

62. Lee, E. H. et al. Association analysis of RGS7BP gene polymorphisms with aspirin intolerance in asthmatic patients. Ann. Allergy. Asthma. Immunol. 106, 292–300 (2011).

63. Wang, B. et al. A block in both early T lymphocyte and natural killer cell development in transgenic mice with highcopy numbers of the human CD3E gene. Proc. Natl. Acad. Sci. 91, 9402–9406 (1994).

64. Luo, M.-L. et al. Amplification and overexpression of CTTN (EMS1) contribute to the metastasis of esophageal squamous cell carcinoma by promoting cell migration and anoikis resistance. Cancer Res. 66, 11690–11699 (2006).

65. Lin, S. et al. Streamlining spatial omics data analysis with Pysodb. Nat. Protoc. 19, 831–895 (2024).

66. Stuart, T. et al. Comprehensive integration of single-cell data. cell 177, 1888–1902 (2019).

67. Gao, C. et al. Iterative single-cell multi-omic integration using online learning. Nat. Biotechnol. 39, 1000–1007 (2021).

68. Welch, J. D. et al. Single-cell multi-omic integration compares and contrasts features of brain cell identity. Cell 177, 1873–1887 (2019).

69. Korsunsky, I. et al. Fast, sensitive and accurate integration of single-cell data with Harmony. Nat. Methods 16, 1289–1296 (2019).

70. Chen, H. et al. Assessment of computational methods for the analysis of single-cell ATAC-seq data. Genome Biol. 20, 1–25 (2019).

71. Lu, M. Y. et al. A visual-language foundation model for computational pathology. Nat. Med. 30, 863–874 (2024).

72. Filiot, A. et al. Scaling Self-Supervised Learning for Histopathology with Masked Image Modeling. medRxiv (2023).

73. Xu, H. et al. A whole-slide foundation model for digital pathology from real-world data. Nature (2024).

74. Antelmi, A. et al. A survey on hypergraph representation learning. ACM Comput. Surv. 56, 1–38 (2023).

75. Kim, S. et al. A survey on hypergraph neural networks: an in-depth and step-by-step guide. in Proceedings of the 30th ACM SIGKDD Conference on Knowledge Discovery and Data Mining 6534–6544 (2024).

76. Feng, Y., You, H., Zhang, Z., Ji, R. & Gao, Y. Hypergraph neural networks. in Proceedings of the AAAI Conference on Artificial Intelligence vol. 33 3558–3565 (2019).

77. Gao, Y., Feng, Y., Ji, S. & Ji, R. HGNN+: General hypergraph neural networks. IEEE Trans. Pattern Anal. Mach. Intell. 45, 3181–3199 (2022).

78. Nair, V. & Hinton, G. E. Rectified linear units improve restricted boltzmann machines. in Proceedings of the 27th International Conference on Machine Learning 807–814 (2010).

79. He, K., Fan, H., Wu, Y., Xie, S. & Girshick, R. Momentum contrast for unsupervised visual representation learning. In Proceedings of the IEEE/CVF Conference on Computer Vision and Pattern Recognition 9729–9738 (2020).

80. Chen, T., Kornblith, S., Norouzi, M. & Hinton, G. A simple framework for contrastive learning of visual representations. in International Conference on Machine Learning 1597–1607 (2020).

81. Velickovic, P. et al. Deep graph infomax. Int. Conf. Learn. Represent. 2, 4 (2019).

82. Wang, Z. et al. NicheTrans: Spatial-aware Cross-omics Translation. bioRxiv 2024–12 (2024).

83. Kingma, D. P. & Ba, J. Adam: A method for stochastic optimization. in International Conference on Learning Representations (2015).

84. Lopez, R., Regier, J., Cole, M. B., Jordan, M. I. & Yosef, N. Deep generative modeling for single-cell transcriptomics. Nat. Methods 15, 1053–1058 (2018).

85. Lin, T.-Y., Goyal, P., Girshick, R. B., He, K. & Dollár, P. Focal Loss for Dense Object Detection. in IEEE International Conference on Computer Vision 2999–3007 (2017).

