## Supplementary Files for "High-Parameter Spatial Multi-Omics through Histology-Anchored Integration"

### Supplementary Text 1

#### Extending SpatialEx+ to multiple consecutive slices

Currently, the SpatialEx+ model only considers the scenario of two consecutive slices. If more consecutive slices are available and their omics information forms a diagonal pattern, it remains unclear how to effectively handle this situation. To address this issue, we proactively propose two potential strategies, referred to as *divide-and-conquer* and *iterative*. Taking three consecutive slices as an example, the two strategies are illustrated correspondingly in Supplementary Fig. 1.

For the *divide-and-conquer* strategy, we enumerated all the paired slices from the dataset and fed them into SpatialEx+ for model training. Having N slices with distinct omics types, we will have (N-1)N/2 separate training steps. The tedious and time-consuming nature of model training is the major drawback.

For the *iterative* strategy, we first randomly selected two slices from the dataset and input them into SpatialEx+ for spatial omics diagonal integration. The resulting completed pair was treated as a pseudo-slice, which was then paired with the next slice and sent into SpatialEx+ to further expand the panel. This iterative process was repeated until the full gene expression matrix was fully completed. For a dataset with N slices, this method requires running SpatialEx+ independently N-1 times, which is more efficient. However, it may lead to error accumulation during the iterative process.

### Supplementary Table 1

| Type | Dataset | Slice | Platform | Modality | Cells | Vars | Mem. (GB) | Time (S) |
|---|---|---|---|---|---|---|---|---|
| H&E to Omics | Human_Breast_Cancer | Rep1 | Xenium | Transcriptomics | 167,780 | 313 | 10.29 | 0.76 |
| | | Rep2 | | | 118,752 | | | |
| | hColon_Non_diseased | S1 | Xenium | Transcriptomics | 134,007 | 325 | 10.08 | 0.73 |
| | | S2 | | | 129,499 | | | |
| | mouse_Colon | S1 | Xenium | Transcriptomics | 107,988 | 379 | 8.68 | 0.61 |
| | | S2 | | | 110,761 | | | |
| | hSkin_Melanoma | S1 | Xenium | Transcriptomics | 43,969 | 282 | 4.01 | 0.29 |
| | | S2 | | | 61,455 | | | |
| Panel Integration | Human_Breast_Cancer | Rep1 | Xenium | Transcriptomics | 167,780 | 150 | 10.32 | 0.84 |
| | | Rep2 | | | 118,752 | 163 | | |
| | Human_Breast_IDC_Big | Rep1 | Xenium | Transcriptomics | 892,966 | 140 | 18.53 | 73.4 |
| | | Rep2 | | | 885,523 | 140 | | |
| Omics Integration | Human_Breast_Cancer | Rep1 | IF | Proteomics | 167,780 | 16 | 11.74 | 0.86 |
| | | Rep2 | Xenium | Transcriptomics | 118,752 | 313 | | |
| | SMA_V11L12-109 | Rep_C1 | Visium | Transcriptomics | 2907 | 1000 | 0.49 | 0.06 |
| | | Rep_B1 | MALDI-MSI | Metabolomics | 3098 | 50 | | |

**Supplementary Table 1: Dataset Statistics.** The table presents key characteristics of each dataset, including sequencing platform, number of cells, number of measured molecules (genes, proteins, or metabolites), and the computational cost of our proposed algorithm. In the "Slice" column, entries prefixed with "Rep" indicate consecutive tissue sections, while others correspond to a single slice artificially divided into two non-overlapping regions.

### Supplementary Figure 1

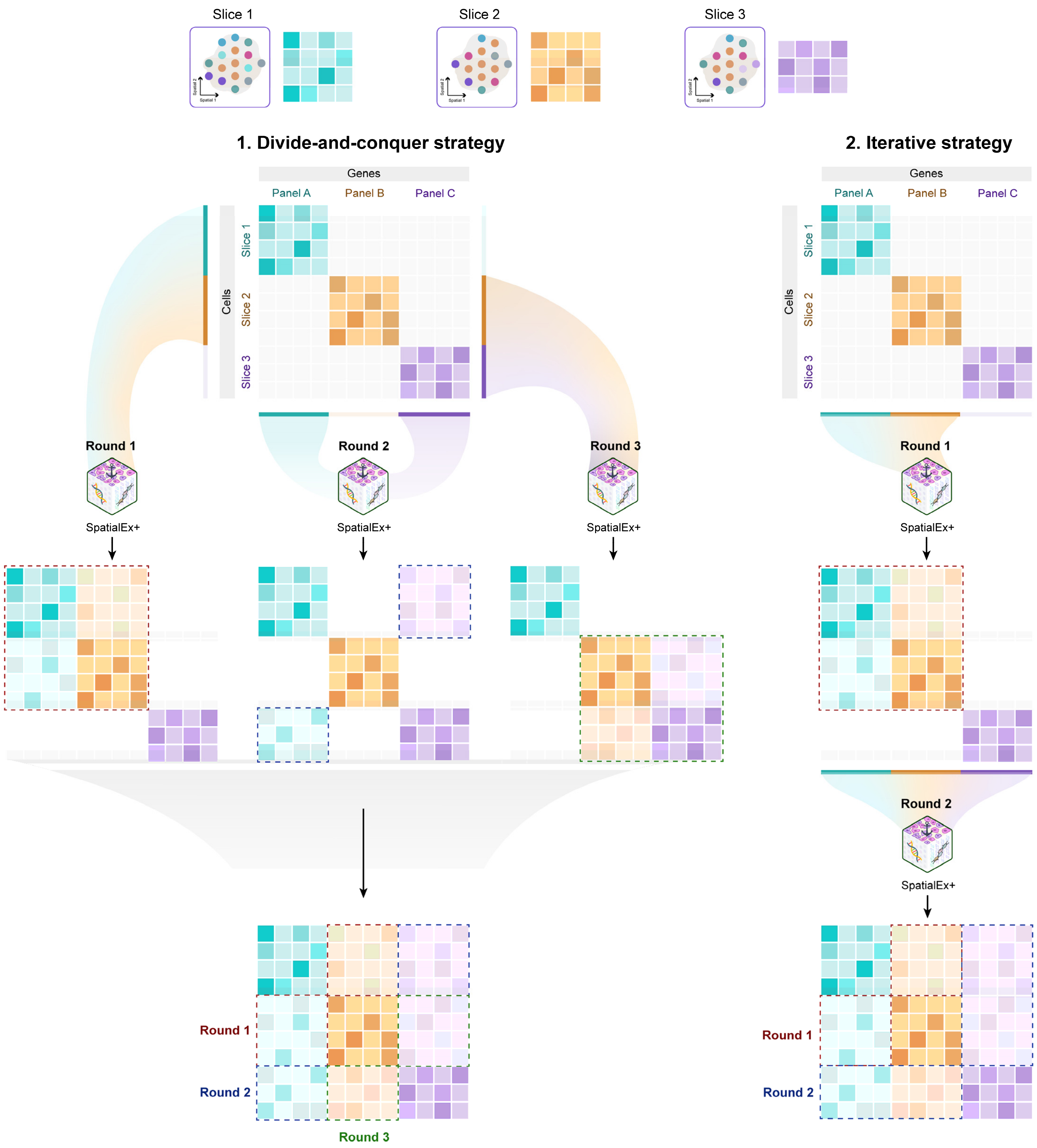

**Supplementary Fig. 1: A schematic illustration of two potential strategies for extending SpatialEx+ to multi-slice scenarios.**

### Supplementary Figure 2

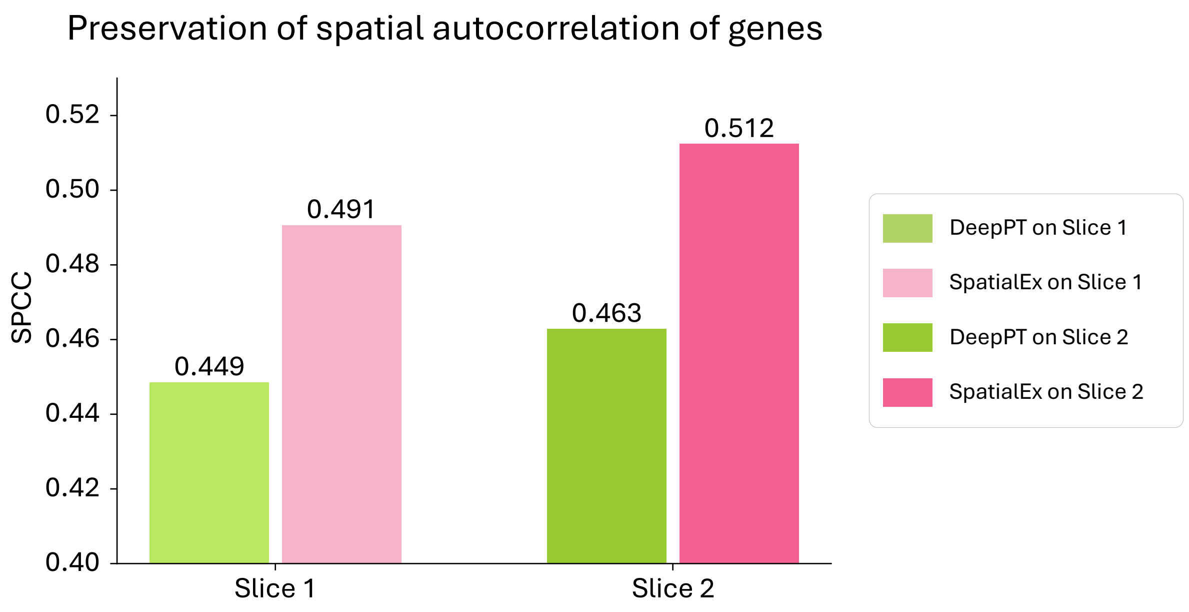

**Supplementary Fig. 2: SPCC comparison of gene expression spatial autocorrelation preservation in H&E-to-omics task.** The similarity in spatial autocorrelation (Moran's *I*) between the measured gene expression and the predictions from DeepPT and SpatialEx is quantified by the Spatial Pearson Correlation Coefficient (SPCC). Bar plots show the SPCC values for both models across two tissue sections, with higher values indicating superior preservation of the original spatial patterns.

### Supplementary Figure 3

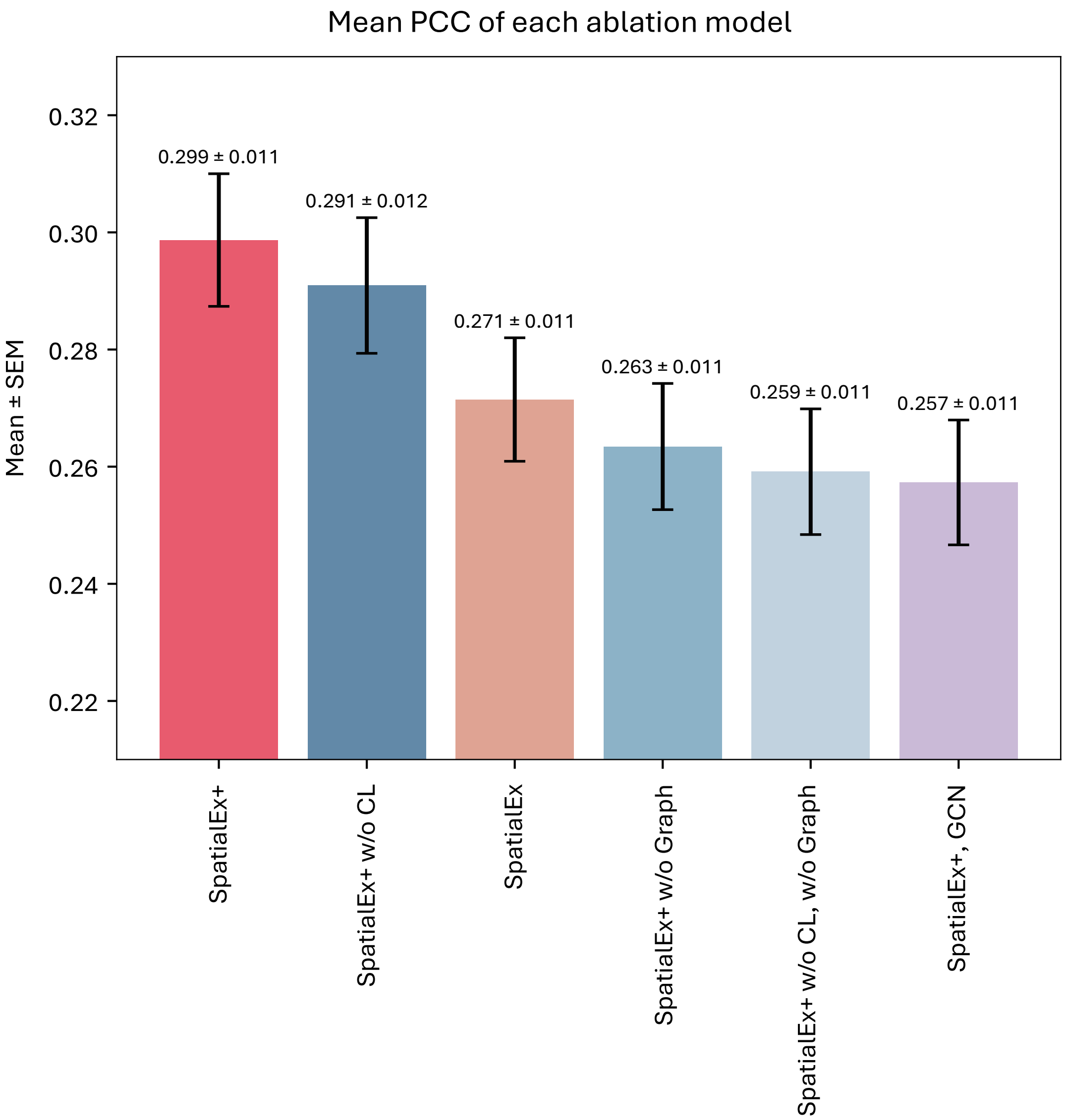

**Supplementary Fig. 3: Ablation study.** The bar plot is about the ablation study of SpatialEx+ on the Human_Breast_Cancer dataset regarding mean PCC. The results demonstrate that each introduced model plays a critical role in the computation and they jointly corporate to achieve the best performance. Error bars represent the standard error of the mean (***n***=313 genes).

### Supplementary Figure 4

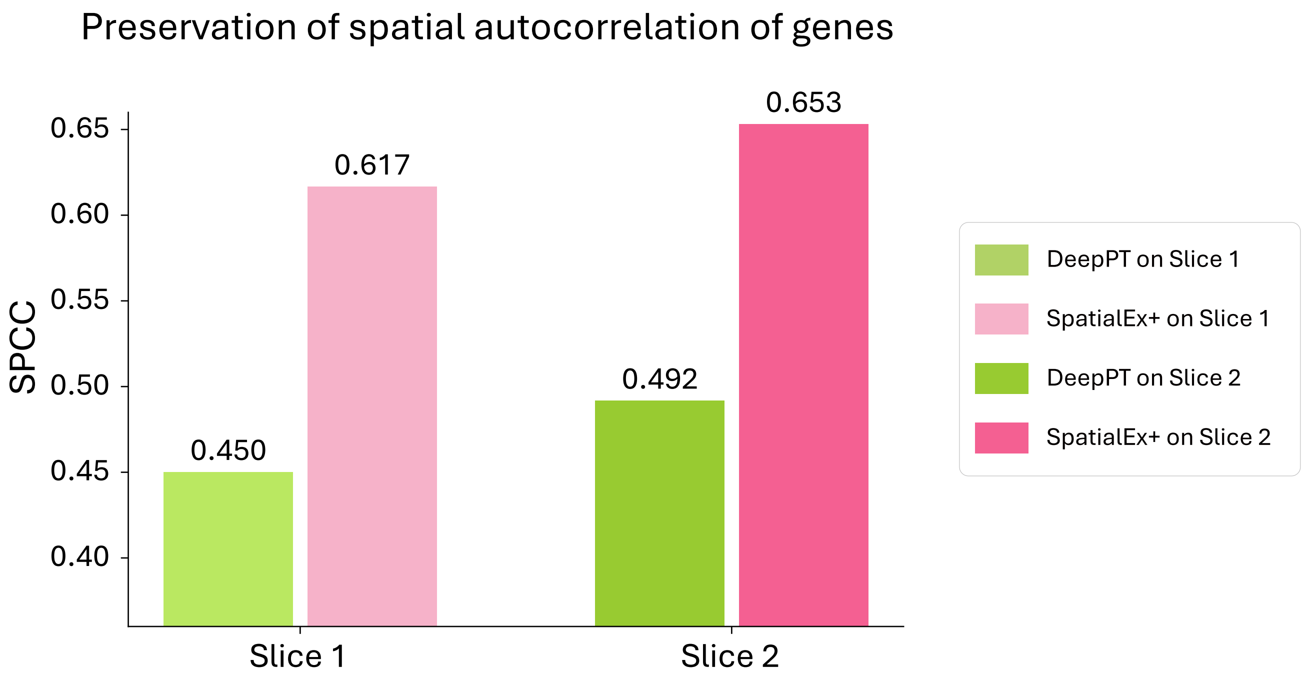

**Supplementary Fig. 4: SPCC comparison of gene expression spatial autocorrelation preservation in panel integration task.** The similarity in spatial autocorrelation (Moran's *I*) between the measured gene expression and the predictions from DeepPT and SpatialEx is quantified by the Spatial Pearson Correlation Coefficient (SPCC). Bar plots show the SPCC values for both models across two tissue sections, with higher values indicating superior preservation of the original spatial patterns.

### Supplementary Figure 5

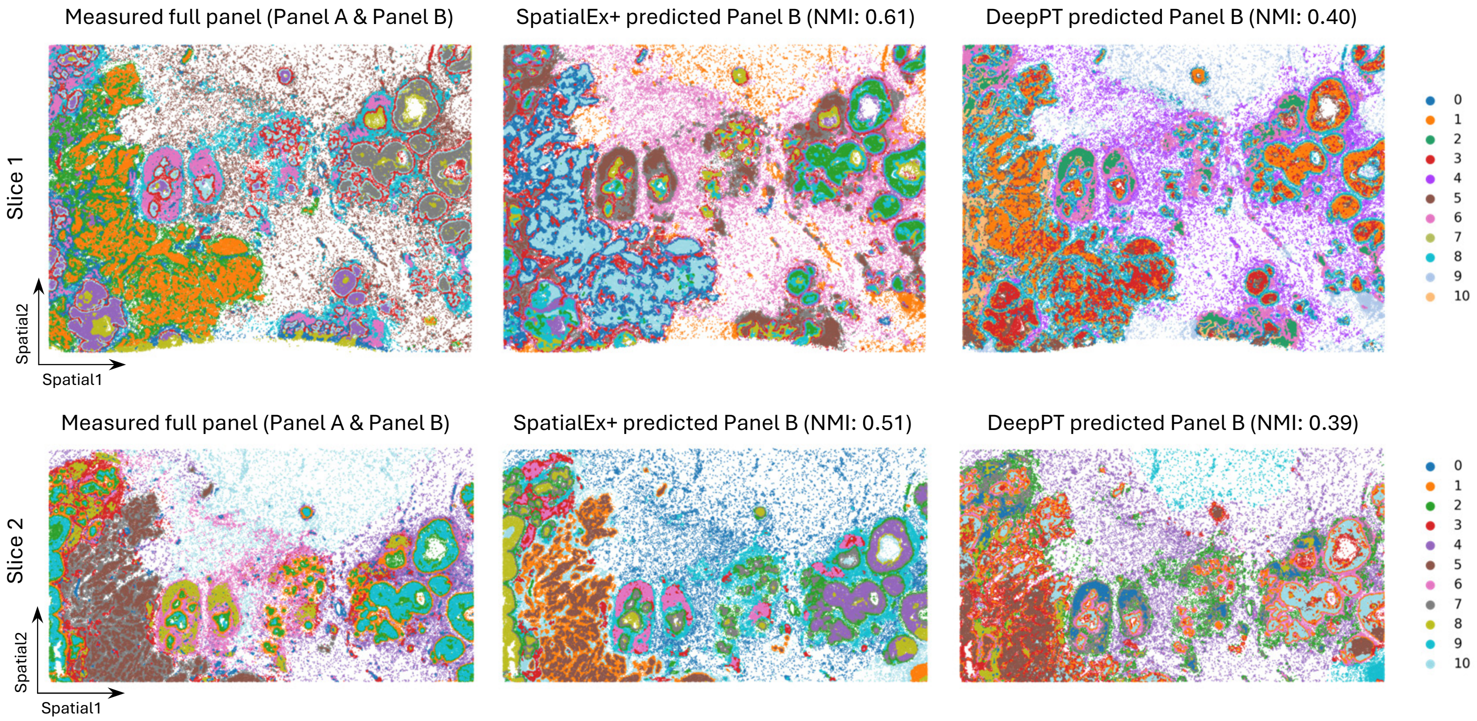

**Supplementary Fig. 5: Spatial domain identification across different conditions on Human_Breast_ Cancer dataset.** Spatial domains derived from experimentally measured full-panel gene expression were used as the ground truth. The identified domains using SpatialEx+ and DeepPT predicted Panel B are illustrated in the second and third columns, respectively. NMIs are utilized as quantitative metrics. Notably, invasive ductal carcinoma (IDC) and ductal carcinoma *in situ* (DCIS) cannot be distinguished using DeepPT predicted Panel B on Slice 1.

### Supplementary Figure 6

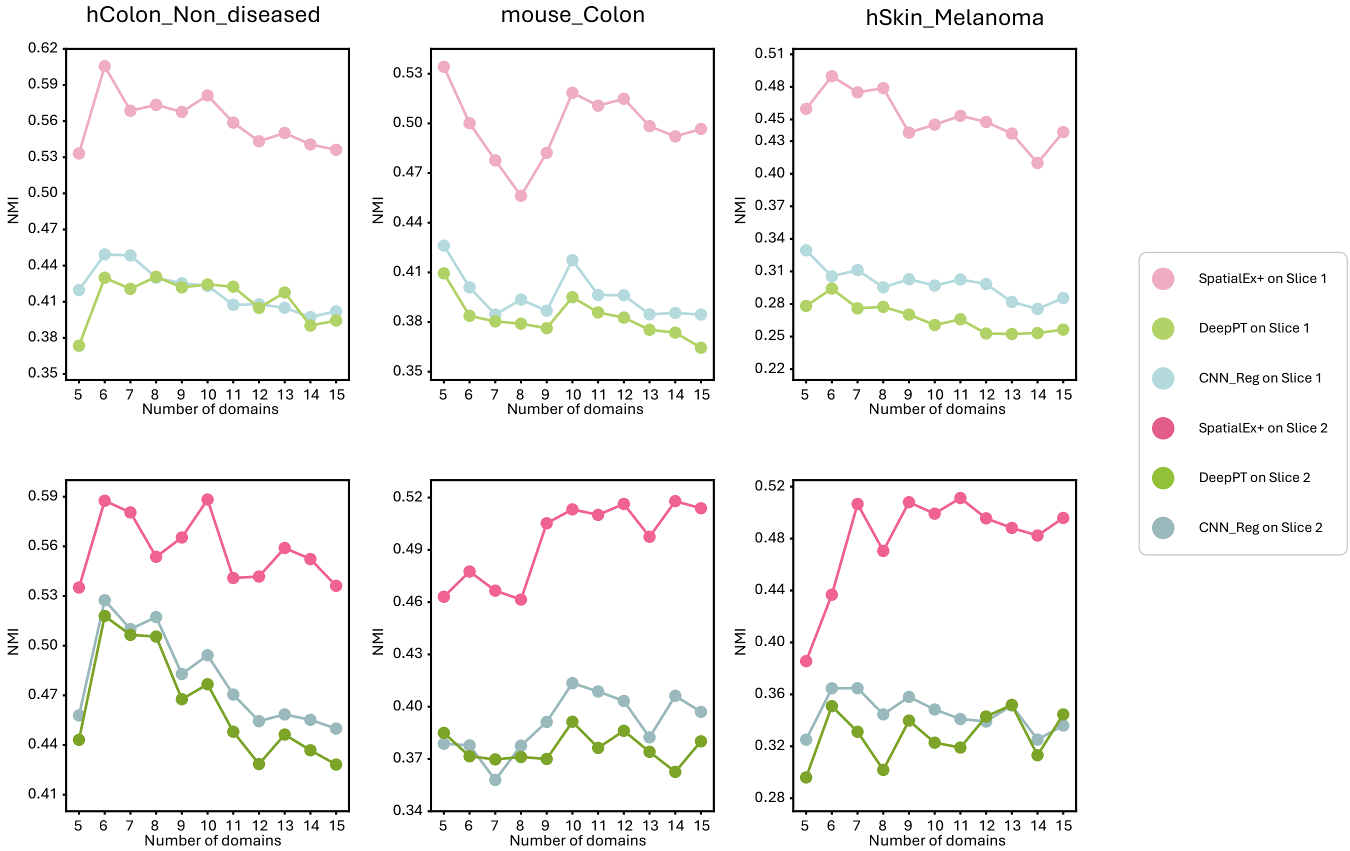

**Supplementary Fig. 6: Line plots of NMI scores across different spatial resolutions.** Spatial domain identification was performed using CellCharter based on measured or predicted gene expression (SpatialEx+, DeepPT, and CNN_Reg. Domains delineated from measured expression were used as ground truth, and NMI was calculated to quantify each method. Across all experimental settings and spatial resolutions, SpatialEx+ consistently produced spatial domains most similar to those derived from measured expression, outperforming both DeepPT and CNN_Reg.

### Supplementary Figure 7

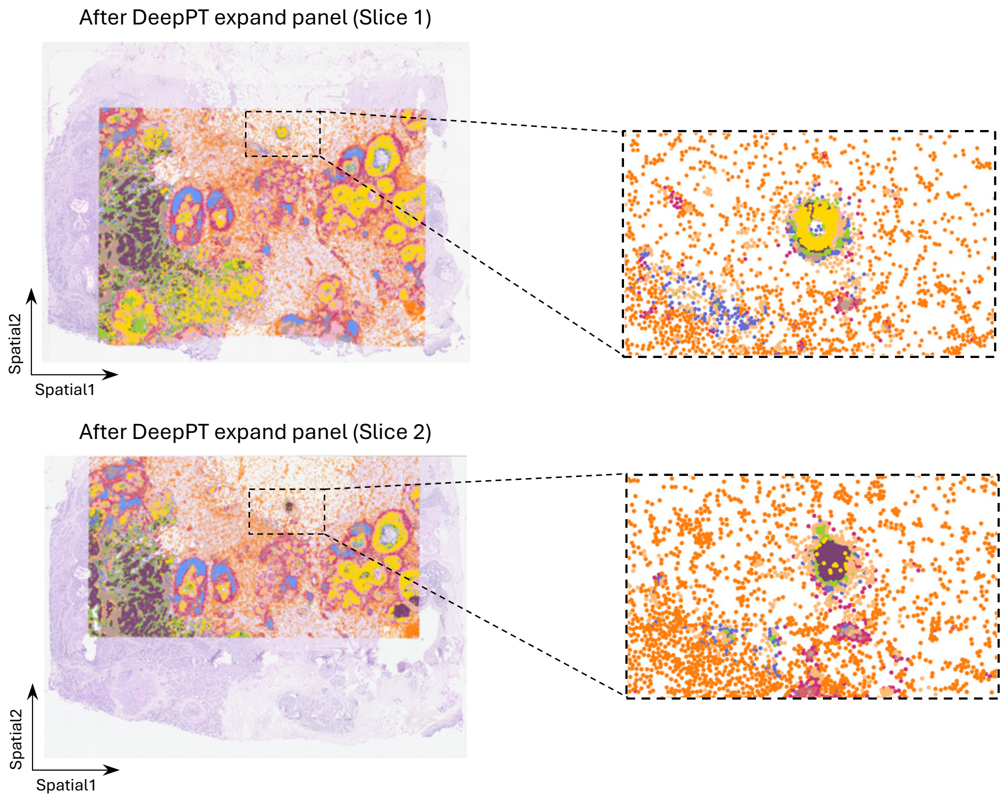

**Supplementary Fig. 7: Integrated spatial domain identification across the two slices using DeepPT expanded full-panel.** Given the inferior prediction capability of DeepPT, the clustering results using the predicted full-panel expression failed to distinguish DCIS and IDC. This limitation also resulted in inconsistent spatial domain assignments for cells at identical anatomical locations, for example in zoomed-in areas.

### Supplementary Figure 8

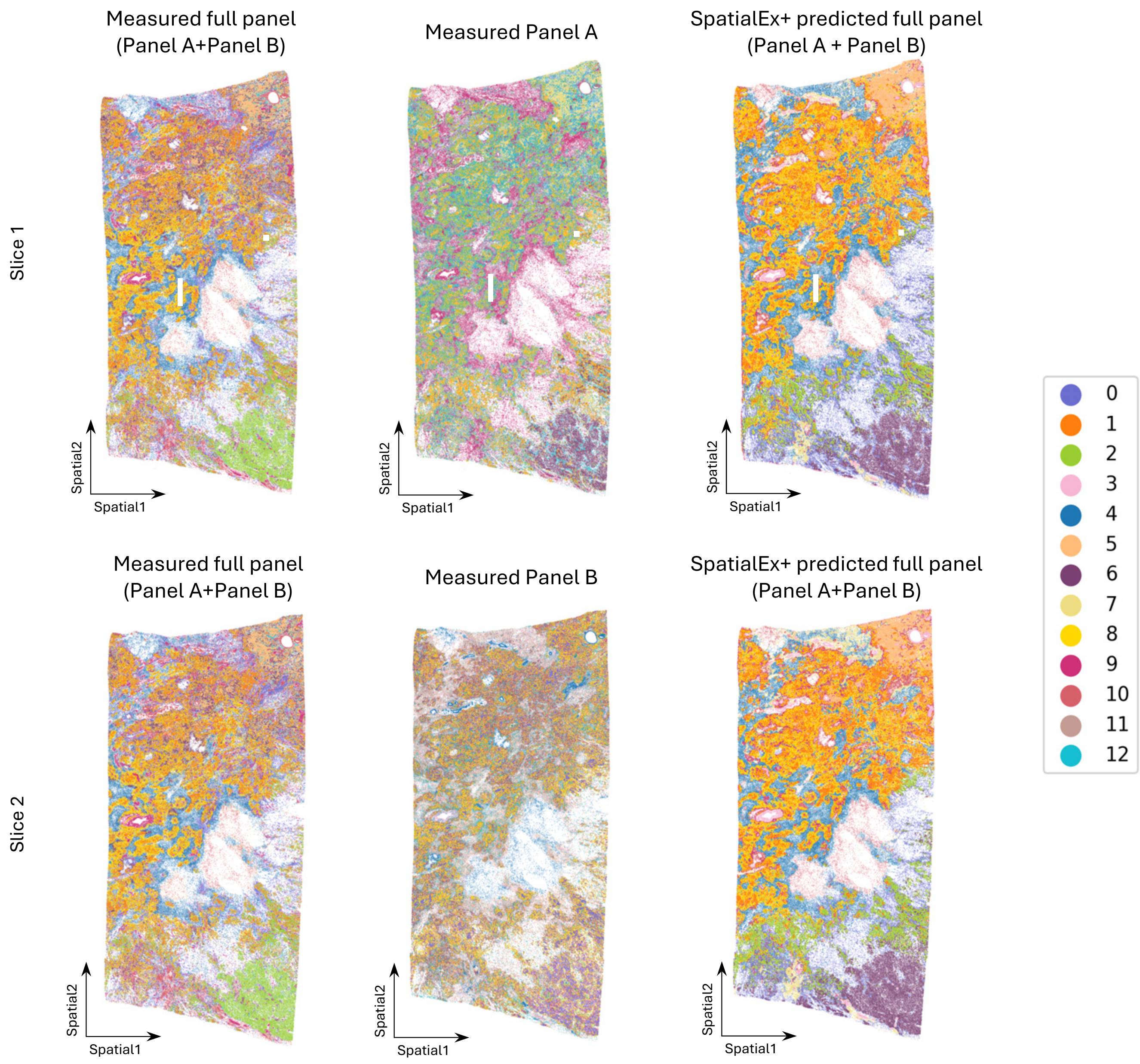

**Supplementary Fig. 8: Spatial domain identification across different conditions on the Human_Breast_IDC_Big dataset.** The first and second rows illustrate the results on Slice 1 and Slice 2, respectively. First column: Joint spatial domain identification on both slices using measured full-panel gene expression data. Second column: Independent spatial domain identification of each slice with measured partial panels (Panel A on Slice 1; Panel B on Slice 2). Third column: Joint spatial domain identification on the two slices using SpatialEx+ predicted full panel (Panel A + Panel B).

#### Supplementary Figure 9

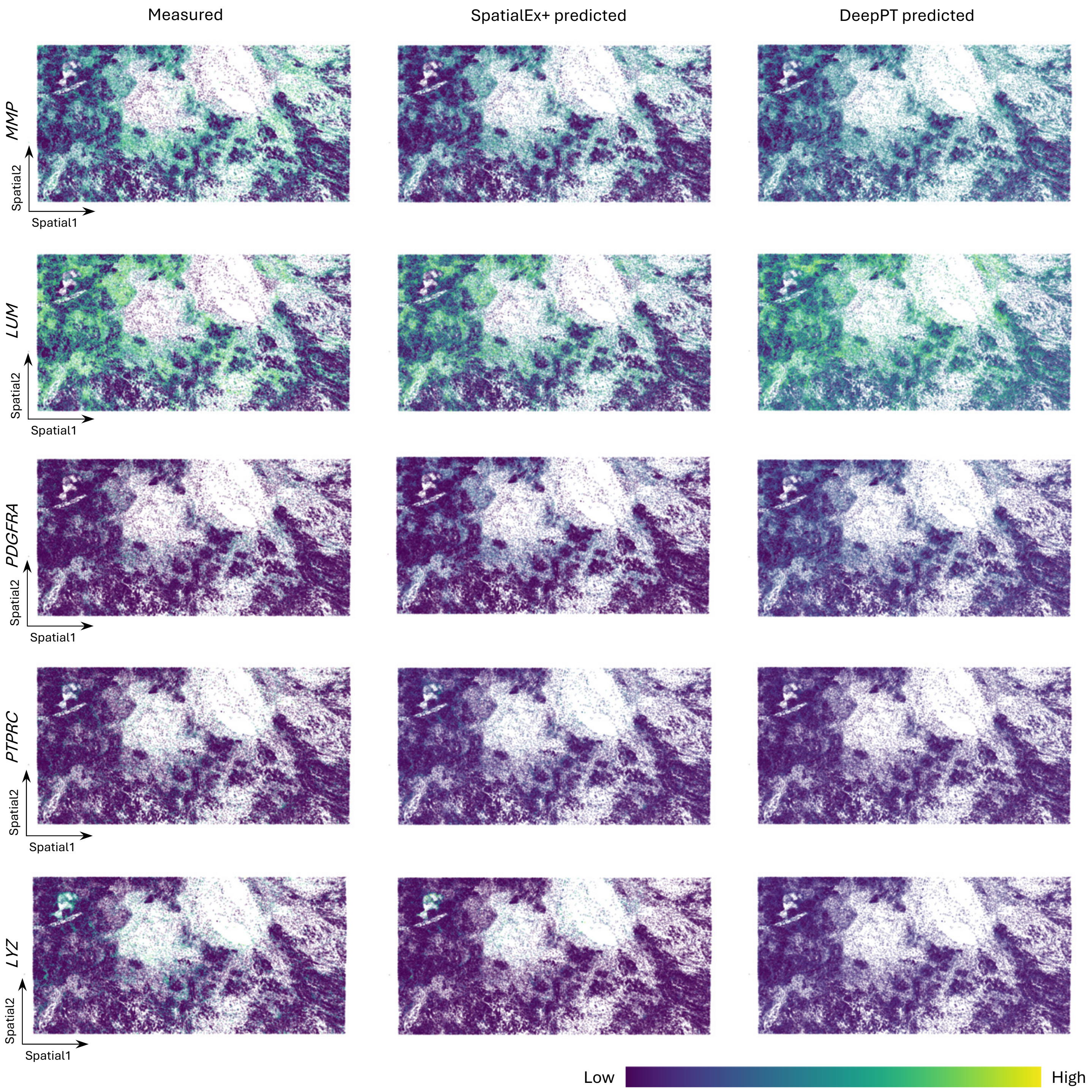

**Supplementary Fig. 9: Visualization of representative DEGs between spatial domain 0&4, and domain 7 on the Human_Breast_IDC_Big Slice 1.** The predictions from SpatialEx+ demonstrate more similar spatial patterns with the measured gene expression, whereas DeepPT failed to recapitulate the transcriptional distinctions between these domains.

### Supplementary Figure 10

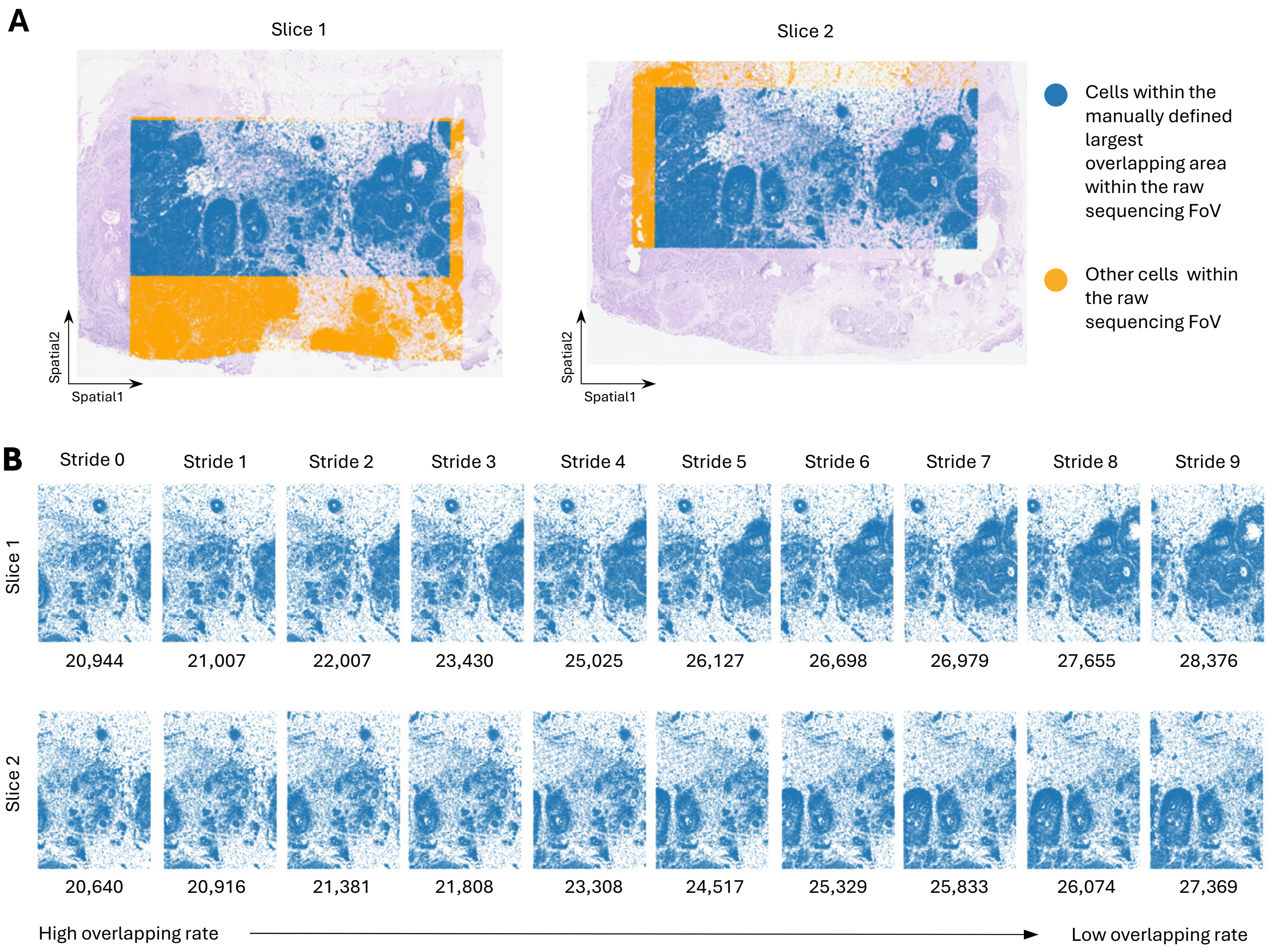

**Supplementary Fig. 10: Training data selection in sliding window experiments. A.** Illustration of the overlapping regions of the two slices regarding original sequencing areas. **B.** Visualization of the cellular distributions along with sliding window striding on. The total number of cells is attached below each window.

### Supplementary Figure 11

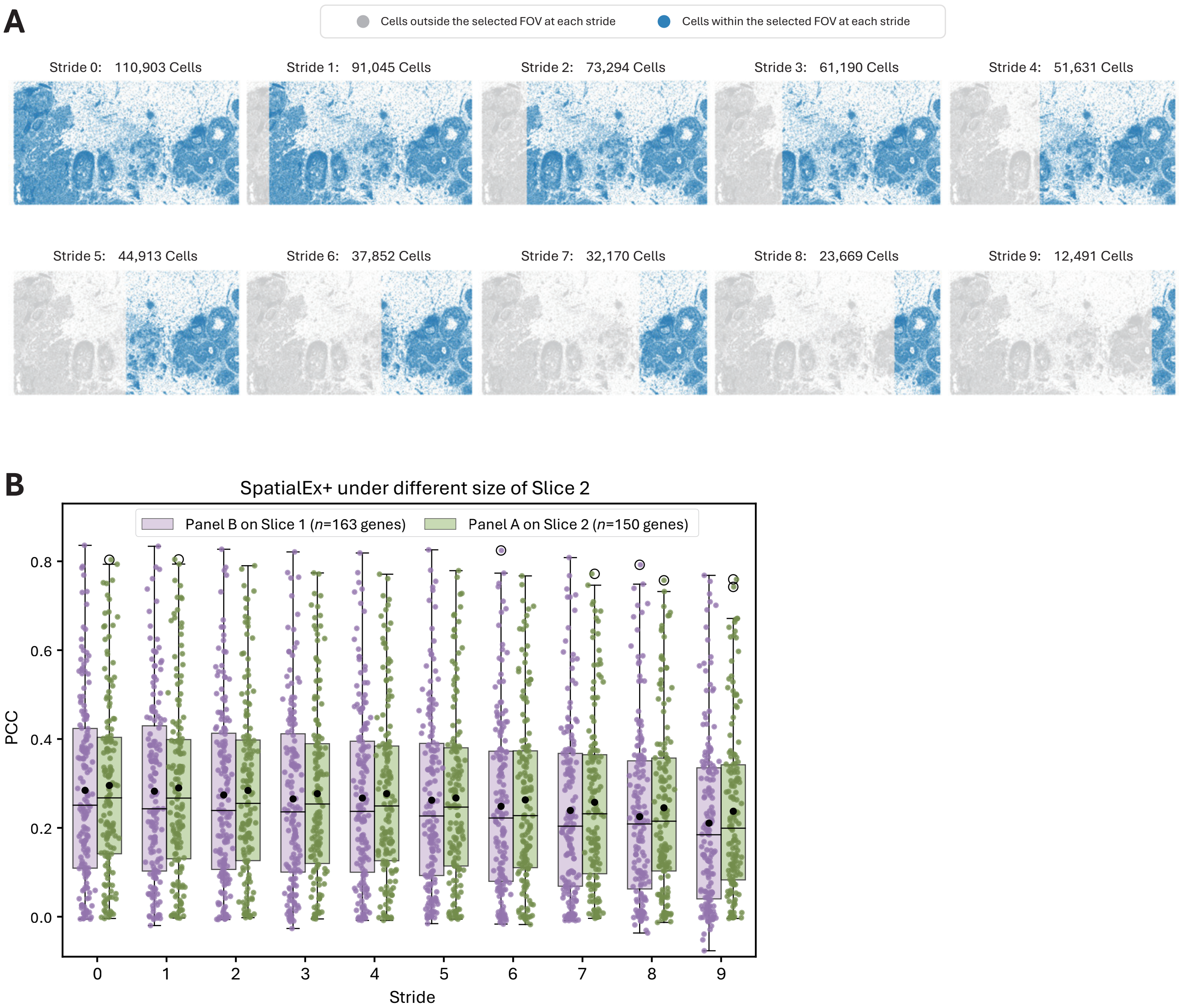

**Supplementary Fig. 11: Performance of SpatialEx+ under varying sizes of Slice 2. A.** Schematic illustration of the number of cells contained in the selected fields of view (FOV) at different stride settings. **B.** A box plot showing the PCC values of SpatialEx+ at each stride, where increasing the stride corresponds to a smaller Slice 2 area. Predictions of Panel A on Slice 2 (***n***=163) genes and Panel B Slice 1 (***n***=150) genes are indicated in green and purple, respectively. Each dot in the scatter represents an individual gene. In the boxplot, the center line, center dot, box limits and whiskers denote the median, mean, upper and lower quartiles and 1.5× interquartile range, respectively.
